# Secondary lymphoid organ endothelial cells prime alloreactive CD4^+^ T cells to trigger acute graft-versus-host disease

**DOI:** 10.1101/2025.10.07.677751

**Authors:** Haroon Shaikh, Pia Wittmann, Moutaz Helal, Michael A.G. Kern, Zahraa Abboud, Juan Gamboa Vargas, Joern Pezoldt, Zeinab Mokhtari, Katja Jarick, Hao Yan, Jörg P.J. Müller, Antoine-Emmanuel Saliba, Angela Riedel, Alma Zernecke, Maike Büttner-Herold, Hermann Einsele, Jochen Huehn, Andreas Beilhack

## Abstract

Donor CD4⁺ T cell priming is a pivotal determinant of acute graft-versus-host disease (aGvHD) after allogeneic hematopoietic cell transplantation (allo-HCT). While professional hematopoietic antigen-presenting cells (APCs) have long been implicated in the pathogenesis of aGvHD, the contribution of non-hematopoietic APCs has remained unclear. Here, we show that naïve alloreactive CD4⁺ T cells initially localize and activate specifically within secondary lymphoid organs (SLOs) before infiltrating target tissues. Using genetic models to selectively ablate MHC class II on endothelial cells (ECs) or hematopoietic cells, we demonstrate that blood endothelial cells (BECs) in SLOs function as APCs, efficiently processing and presenting antigen to prime donor CD4⁺ T cells. Deletion of MHC class II (MHCII) specifically in ECs substantially attenuates T cell activation and protects mice from lethal aGvHD, whereas selective deletion of MHCII in lymphatic ECs has no effect. Likewise, selective deletion of MHCII in hematopoietic cells also protects mice against aGvHD, suggesting that both cell types contribute to pathogenic allogeneic T cells activation after allo-HCT. Mechanistically, IL-12/IFNγ signaling upregulates MHC class II expression on BECs. These findings identify BECs in SLOs as initiators of alloreactive CD4⁺ T cell responses and highlight a potential target for preventing aGvHD.

**Highlights:**

- Blood endothelial cells in secondary lymphoid organs prime naïve CD4⁺ T cells to trigger acute GvHD.
- T cell activation occurs exclusively in secondary lymphoid organs before tissue infiltration.
- Expression of MHC class II only on endothelial cells is sufficient to drive lethal GvHD, independent of other antigen presenting cells.
- Regulation of MHC class II expression in blood endothelial cells by IL-12/IFNγ offers the potential for new therapeutic targets and corroborates findings for existing therapeutics.

## Introduction

Allogeneic hematopoietic cell transplantation (allo-HCT) remains a potentially curative therapy for hematologic malignancies, yet its efficacy is limited by acute graft-versus-host disease (aGvHD) (Zeiser and Blazar). aGvHD is a life-threatening complication of allo-HCT that results from donor T cell recognition of “self” allo-antigens presented on major histocompatibility complex (MHC) molecules (Duffner et al.; Shlomchik et al.). While donor CD8⁺ T cells can recognize MHC class I directly, initiation of donor CD4⁺ T cell responses depends on presentation by MHC class II (MHCII)-expressing antigen-presenting cells (APCs) (Beilhack et al.). Identifying the precise location and cellular source of CD4⁺ T cell priming is therefore essential for understanding how alloreactivity is initiated.

Professional hematopoietic APCs – dendritic cells (DCs), macrophages, and B cells – are classically considered the main initiators of MHCII–restricted responses (Unanue) and are thought to be the primary instigators of aGvHD.

However, several landmark studies have shown that ablating MHCII expression on hematopoietic APCs does not fully prevent aGvHD (Koyama et al.; Li et al.; Toubai et al.), suggesting the possibility that non-hematopoietic cells might also present antigen and prime pathogenic CD4⁺ T cells. Indeed, non-hematopoietic lineage cells, including endothelial, epithelial, and stromal cells, express MHCII in response to IFN-γ stimulation and contribute to immune responses (Kambayashi and Laufer; Londei et al.). Yet, the physiological significance of such expression in priming pathogenic CD4⁺ T cells remains controversial and incompletely understood.

A central clue in the context of allo-HCT lies in the migration pattern of donor T cells. Naïve donor CD4⁺ T cells first accumulate in secondary lymphoid organs (SLOs) and only later infiltrate target tissues such as the gut, liver, and skin (Beilhack et al.; Beilhack et al.). Because naïve T cells require priming before acquiring expression of tissue-homing receptors (Mora et al.), their early localization to SLOs strongly suggests that the relevant non-hematopoietic APCs reside there rather than in peripheral target organs. As such, whereas epithelial cells in the gut or skin have been implicated in amplifying effector responses in aGvHD (Koyama et al.), they are unlikely to initiate the first CD4⁺ T cell activation step.

Lymph nodes (LNs), Peyer’s patches (PPs) and the spleen are pivotal SLOs that orchestrate adaptive immune responses via specialized stromal and hematopoietic niches (Bajenoff et al.; Junt et al.; Sixt et al.; Warnock et al.). Comprising approximately 98% leukocytes and 2% non-hematopoietic stromal cells, LNs integrate mesenchymal and endothelial components to provide structural support and immunoregulatory cues to T and B cells (Krishnamurty and Turley). The primary stromal cells in LNs include fibroblastic reticular cells, blood endothelial cells (BECs), and lymphatic endothelial cells (LECs), all of which express MHCII (Card et al.; Jalkanen and Salmi; Onder et al.; Perez-Shibayama et al.; Potente and Makinen). However, *Ccl19*^+^ fibroblastic reticular cells, which are the dominant cell type in LN T cell zones, do not prime allogeneic T cells during aGvHD. Indeed, selective absence of MHCII on these cells (MHCII^ΔCcl19^) exacerbated aGvHD due to reduced expansion of regulatory T cells (Shaikh et al.). LECs also promote tolerance during homeostasis and in inflammatory conditions (Cohen et al.; Dubrot et al.; Gkountidi et al.; Rouhani et al.; Tewalt et al.). In contrast, the role of BECs during aGvHD initiation and naïve allogeneic CD4⁺ T cell priming has been neglected, although they play a prime role in triggering trans-endothelial migration and activation of circulating effector memory CD4^+^ T cells (Pober et al.).

This unresolved question has critical implications. If SLO BECs function as non-hematopoietic APCs during allo-HCT, they would represent a previously unrecognized entry point into the allogeneic cascade and a potential therapeutic target. Clarifying this issue is therefore essential not only to resolve prior controversies, but also to enable rational design of strategies that selectively prevent pathogenic donor T cell priming without compromising beneficial graft-versus-leukemia effects.

Here, we identify SLO BECs as central non-hematopoietic APCs that present alloantigen to naïve donor CD4⁺ T cells and initiate lethal aGvHD. These findings redefine the cellular framework of alloimmune priming and highlight the vascular endothelium of SLOs as a critical regulator of transplant immunity.

## Results

### Allogeneic CD4^⁺^ T cells are activated and expand within SLOs before infiltrating the gut in aGvHD initiation

As a first step to identifying which non-hematopoietic APCs initiate allogeneic CD4⁺ T cell responses during aGvHD, we first asked where donor T cells encounter alloantigen and undergo priming. Defining this site is critical, as only by mapping the earliest stages of antigen engagement, clonal expansion, and effector programming is it possible to distinguish true priming from downstream infiltration and to determine which non-hematopoietic APCs in which tissues drive initial immune activation.

In a major MHC-mismatched allo-HCT model (FVB➔C57BL/6) (**Figure 1A**), CD4⁺ T cells rapidly homed to SLOs, including PPs, mesenteric LNs (mLNs), and spleen, within six hours after transplantation (**Figure S1A**). Bioluminescence imaging revealed steadily increasing signals in these tissues over the first 76 hours (**Figure 1B**), whereas the small intestinal lamina propria remained devoid of donor T cells (**Figure 1B, Figure S1B**). By day 4, donor T cells started to appear outside of SLOs, with progressive infiltration of the small intestinal lamina propria peaking around day 6 (**Figure 1B**), consistent with earlier findings (Bauerlein et al.; Beilhack et al.; Beilhack et al.; Brede et al.). Several reports have suggested that CD4⁺ donor T cells can be primed outside of SLOs after allo-HCT; for example, priming in intestinal tissues has been attributed to antigen presentation by intestinal myofibroblasts (Koyama and Hill; Koyama et al.) and intestinal epithelial cells (Koyama and Hill; Koyama et al.; Koyama et al.). Given that bioluminescence imaging lacks the cellular resolution and spatial precision to identify the location of T cell priming, we set out to test whether donor CD4⁺ T cell priming can occur in the intestinal lamina propria.

**Figure 1:** Alloreactive CD4⁺ T cells are primed in secondary lymphoid organs during the initiation of aGvHD **(A)** Experimental design: Myeloablatively irradiated (9 Gy) B6.WT recipients were transplanted with 5x10^6^ TCD BM and 5x10^6^ CFSE labelled luc^+^ CD4^+^ T cells (i.v.) from FVB and FVB.L2G85 donors, respectively. **(B)** *Ex vivo* bioluminescence imaging (BLI) of the gastrointestinal tract and spleen at days 1-6 after allo-HCT. Signals accumulated in secondary lymphoid organs but not in peripheral tissues during the first 3 days. Data are representative of 2 replicate experiments (n = 3-4). **(C)** 3D light sheet fluorescence microscopy (LSFM) of ileum at days +1–3 after allo-HCT. CD31⁺ blood vessels (red), donor CD45.1⁺ CD4⁺ T cells (green), and tissue autofluorescence (grey). Scale bars: 300 μm. Donor T cells localized almost exclusively within Peyer’s patches and in direct apposition to vasculature using a 0Lμm distance threshold. Quantification performed with Imaris. Representative of 2 experiments (n = 3-4). **(D)** 3D-Swiss roll LSFM of ileum at day +3 (top panel) and +6 (lower panel) of allo-HCT. Tissue autofluorescence (grey), with CD45.1 with AF647 (green) and CD31 biotin and subsequently with streptavidin AF750 (red). **(E)** Confocal images of Peyer’s patches and adjacent small intestinal lamina propria (SILP) at days +3 and +6 of allo-HCT. Laminin (magenta), donor CD45.1⁺ T cells (green), nuclei (grey). Infiltration of donor T cells into the SILP was absent at day +3 and evident by day +6. Scale bar: 100 μm. Representative of 2 experiments (n = 2). **(F)** Myeloablatively irradiated (9 Gy) C57BL/6 and B6.MHCII^Δ^ mice were i.v. transplanted with 5x10^6^ TCD BM and 5x10^6^ CD4^+^ T cells from FVB. Expression of CD44, integrin α4β7 and CCR9 in MFI histogram on donor T cells (CD90.1^+^CD4^+^) day 3 of allo-HCT in and quantification in MFI fold change in the mesenteric lymph nodes and Peyer’s patches. Data is representative of 1 experiment (n = 3). **(G)** Correlation heatmap of protein fold changes across C57BL/6 and B6.MHCII^Δ^ mouse in Peyer’s patches at day +3 of allo-HCT. Pearson correlation coefficients (r) were calculated between fold changes of CD62L, CD44, α4β7, and CCR9 expression levels across 6 samples (3x C57BL/6 and 3x B6.MHCII^Δ^). The heatmap displays pairwise correlations, with hierarchical clustering applied to both rows and columns. Color scale represents correlation values from –1 (strong negative correlation) to +1 (strong positive correlation), with white indicating no correlation (r = 0). Heatmaps were generated using the pheatmap R package. **(H)** Flow cytometry analysis of donor CD4^+^ T cells (CD45.1^+^CD4^+^) for CFSE dilution from spleen, mLNs and small intestine at day 3 of allo-HCT and quantification. Data are representative of 3 replicate experiments (n = 3-4). Statistical analysis by two-way ANOVA with Sidak’s multiple comparison test (F) and ordinary one-way ANOVA, adjusted for multiple comparisons with Tukey’s multiple-comparison test (H), (Mean± SD); \**p <* 0.05, \*\**p <* 0.01. i.v.: intra-venous, luc: luciferase, GIT: gastrointestinal tract, LSFM: Light sheet fluorescent microscopy, CFSE: Carboxyfluorescein succinimidyl ester, SILP: small intestine lamina propria.

To map the location of donor T cells in high resolution across large tissue volumes, we used three-dimensional light sheet fluorescence microscopy (3D-LSFM). From days 1–3, CD45.1⁺ donor CD4⁺ T cells localized almost exclusively within SLOs such as PPs, with very few cells detectable in the small intestinal lamina propria (**Figure 1C and Figure S1B**). Strikingly, nearly all donor T cells associated with small intestinal lamina propria resided intravascularly (**Figure S1C**), indicating circulating cells rather than activation. Confocal imaging and whole-tissue “Swiss roll” LSFM (Mueller et al.) of the ileum a region highly prone to barrier disruption and immune infiltration (Hulsdunker et al.) confirmed that extravascular donor CD4^+^ T cells remained absent from the lamina propria parenchyma through day 3, but were abundant by day 6 (**Figure 1D, E** and **Figure S1D, E**).

To test whether donor CD4⁺ T cells could acquire expression of gut-homing receptors independently of alloantigen recognition, we transferred donor T cells into recipient mice lacking MHCII. In this setting, donor T cells in PPs and mLNs retained a naïve CD62L^hi^CD44^lo^ phenotype and failed to induce α4β7 and CCR9 at day 3 after transplantation (**Figure 1F, G)**, demonstrating that priming is indispensable for activation and expression of gut-homing receptors. Consistently, analysis of donor CD4^+^ T cell proliferation revealed robust clonal expansion in spleen and mesenteric LNs at day 3, whereas the few T cells found in the small intestinal lamina propria at this time point had fully diluted the CFSE label, consistent with prior activation in SLOs (**Figure 1H**).

Taken together, these data establish that CD4⁺ T cells are activated and expand within SLOs before infiltrating the gut. Priming in the small intestinal lamina propria appears to be an extremely rare event, if it occurs at all, suggesting that this tissue has little to no contribution to early immune activation during aGvHD.

### Lymph node endothelial cells express MHC class II and process exogenous antigens

We next investigated which non-hematopoietic APCs in SLOs are capable of initiating alloreactive priming. To define the capacity of different populations of LN endothelial cells (ECs) to present antigen, we re-analyzed previously generated single-cell RNA-sequencing (scRNA-seq) data of mesenteric and peripheral LNs of adult BALB/c mice (Pezoldt et al.). Among CD45⁻CD24⁻ stromal cells, ECs were selected based on *Pecam1* (CD31) expression, and using unbiased uniform manifold approximation and projection (UMAP) clustering, we identified transcriptionally distinct EC subsets with characteristic molecular (**Figure 2A, B**). Arterial BECs expressed *Esm1, Cd146*, *Cd36*, *Cxcr4*, *Ets2, and Itga1*, whereas venous BECs expressed *Ackr1, Glycam1*, *Madcam1*, *Lrg1*, and *Fut7*. LECs were defined by *Prox1 and Pdpn* expression and further stratified into ceiling LECs of the sub-capsular sinus (*Ackr4*, *Cd36*, *Anxa2*, and *Bmx*) and combined floor and medullary LECs (*Glycam1*, *Ccl20*, *Coch*, *Marco*, and *Itga2b*) (**Figure 2C** and **Figure S2A**).

**Figure 2:** LN ECs express MHCII and co-stimulatory receptors. (A) Gating of LN non-hematopoietic cells: Viable singlets were pre-gated on CD45^-^CD24^-^ and gated for LECs as CD31^+^gp38^+^ whereas BECs as CD31^+^gp38^+^ population. (B) In scRNA seq dataset, ECs were identified as *Pecam*^+^ and were selected for further downstream analysis. Data shown is pooled from 2 mLNs (sample 1 = 577 cells, sample 2 = 600 cells) and 2 pLNs (sample 1 = 343 cells, sample 2 = 493 cells) data sets. UMAP plot of merged mLNs ECs and pLNs ECs showing cluster segregation. (C) Expression of subset defining DEGs across ECs on UMAP plot. (D) Heatmap of expression of genes involved in MHCII-mediated antigen presentation on 4 identified clusters of LN ECs. (E) Flow cytometry expression analysis of CD80, CD86 and MHCII shown in histograms on LEC and BEC at steady-state and 24 hours post TBI and quantification in MFI fold change. Data are representative of 3 replicate experiments (n = 3-4). (F) DQ-OVA incubated with FACS sorted LEC and BEC from C57BL/6 mouse and cultured for 3 h at 4°C or 37°C, followed by analysis of processed DQ-OVA under the blue laser (488 nm), BL-1. Data are representative of 3 replicate experiments (n = 3-4). (G) 3D-LSFM stack of mLNs: Blood vessels were stained with *in vivo* primary CD31 biotin and subsequently with streptavidin AF750 (red) and donor CD4^+^ T cells with CD45.1 with AF647 (green) at day +1, +2 and +3 allo-HCT. Scale bar: 40 µm. Data are representative of 2 replicate experiments (n = 3-4). Statistical analysis by unpaired non-parametric Mann-Whitney test (E), one-way ANOVA, adjusted for multiple comparisons with Tukey’s multiple-comparison test (G), (Mean± SD); \**p <* 0.05, \*\**p <* 0.01 and \*\*\*\**p* < 0.0001. mLNs: mesenteric lymph nodes, pLNs: peripheral lymph nodes, ECs: endothelial cells, LECs: lymphatic endothelial cells, BECs: blood endothelial cells, scRNA seq: single cell RNA sequencing, DEGs, differentially expressed gene, UMAP: Uniform manifold approximation and projection for dimension reduction, MFI: mean fluorescent intensity, TBI: Total body irradiation, DQ-OVA: BODIPY-conjugated OVA.

We next interrogated the expression of genes required for MHCII antigen processing and presentation. All EC subsets expressed the lysosomal markers *Lamp1* and *Lamp2*, invariant chain (*li*/*Cd74*), the proteases cathepsin L (*Ctsl*) and cathepsin S (*Ctss*), and classical MHCII genes (*H2-Aa*, *H2-Ab1*, *H2-Eb1*), and all subsets lacked expression of *H2-Eb2* (**Figure 2D**). Other proteases were variably expressed, with cathepsin H (*Ctsh*) enriched mostly in medullary and floor LECs and cathepsin C (*Ctsc*) expressed in a subset of both venous and arterial BECs. All EC subsets expressed a broad array of IFNγ response genes (*Ifngr1/2*, *Stat1*, *Irf1*, *Socs1*), consistent with the potential for IFNγ-inducible MHCII expression (Muhlethaler-Mottet et al.), whereas Toll-like receptors expression was negligible (**Figure S2B**).

Notably, both LECs and BECs expressed the costimulatory molecules CD80 and CD86 at steady-state, in contrast to what has been shown for ECs from non-lymphoid tissues (Amersfoort et al.). Following total body irradiation (TBI), LECs upregulated CD80 and CD86 but downregulated MHCII (**Figure 2E**), whereas BECs upregulated CD80 and CD86, albeit at lower levels, whereas MHCII expression was unchanged. The decrease in MHC class II expression in LECs might reflect the loss of radio-sensitive DCs, which normally transfer MHCII to LN stromal cells under homeostasis and inflammation (Dubrot et al.; Shaikh et al.). To directly test antigen-processing capacity, we cultured FACS-purified LN-derived LECs and BECs with labeled DQ-OVA. Both subsets efficiently degraded the probe, confirming that SLO ECs can internalize and process exogenous antigens at steady-state (**Figure 2F**).

Finally, we examined whether donor CD4⁺ T cells physically interact with SLO ECs during early allo-HCT. 3D-LSFM imaging of mesenteric LNs revealed that at day +1 most donor CD4⁺ T cells resided within 2–25 µm of CD31⁺ blood vessels (**Figure 2G**). By days +2–3, T cells remained associated with vessels but were dispersed further from the endothelium, consistent with a transition from perivascular docking to interstitial migration (**Figure 2G** and **Figure S2C**).

Together, these findings show that LN BECs and LECs express MHCII (both at steady-state and after TBI), costimulatory ligands, and genes encoding components of the antigen processing machinery, positioning them as candidate non-hematopoietic APCs capable of priming alloreactive CD4⁺ T cells during aGvHD initiation.

### Blood-endothelial cells drive alloimmune priming via MHCII

Having established that SLO ECs express MHCII and can process antigen, we next asked whether they functionally prime donor CD4⁺ T cells during aGvHD. We first validated pan-endothelial identity in our scRNA-seq dataset. SLO ECs expressed canonical markers such as VE-cadherin (*Cdh5*) (**Figure 3A**) and TEK tyrosine kinase (*Tie2*) (**Figure S2A**). Although *Tie2* is also expressed by specialized macrophage subsets (Chen et al.; Venneri et al.; Welford et al.), previous research has shown that macrophages primarily regulate aGvHD in similar models (Flamann et al.; Hashimoto et al.; Le et al.), suggesting that ECs are more likely to be the relevant non-hematopoietic APCs *in vivo*.

**Figure 3:** ECs upregulate MHCII post allo-HCT and ECs specific MHCII knockout results in partial aGvHD abrogation. **(A)** Expression of *Cdh5* across LN ECs evaluated by scRNA-seq on UMAP plot. **(B)** Myeloablatively irradiated (9 Gy) B6.*H2-Ab1*^fl^ and B6.MHCII^ΔCdh5^ mice were i.v. transplanted with 5x10^6^ T cell-depleted (TCD) BM and 6x10^5^ CD4^+^ T cells from FVB. Survival by Kaplan-Meier estimates, combined from 3 replicate experiments. **(C)** Expression of *Prox1* across LN ECs evaluated by scRNA-seq on UMAP plot. **(D)** Myeloablatively irradiated (9 Gy) B6.*H2-Ab1*^fl^ and B6.MHCII^ΔProx1^ mice were i.v. transplanted with 5x10^6^ T cell-depleted (TCD) BM and 6x10^5^ CD4^+^ T cells from FVB. Survival by Kaplan-Meier estimates, combined from 2 replicate experiments. **(E)** Expression of invariant chain (li), H2-M and CLIP:I-A^b^ complex on LN blood endothelial cells (Zombie^-^CD45^-^gp38^-^CD31^+^) plotted against MHC-II and displayed as contour plots (n = 3). **(F)** Blood endothelial cells from the pooled lymph nodes (VIVID^-^CD45^-^CD24^-^gp38^-^CD31^+^) from B6.WT mice at steady-state, day 3 post-TBI and day 3 post-allo-HCT were sort purified and processed for RNA-seq gene expression analysis (n = 3). First and second principal component projections (PCA). **(G)** Clustered heatmaps of the genes involved in MHC class II and interferon signalling that were significantly differentially increased (< 0.05) between groups of samples analyzed, plotted as expression Z-score. **(H)** Expression of IFNγ in B6.IFNγ-eYFP mouse at steady-state and day 2 post-TBI on Mφ (CD45^+^CD11c^+^), CD45^+^CD11c^+^CD11b^+^ cells, granulocytes (CD45^+^CD11c^+^CD11b^+^), B cells (CD45^+^CD19^+^), T cells (CD45^+^CD3ε^+^) and ECs (CD45^-^CD31^+^) and quantification in MFI. **(I)** Expression of IL-12/IL-23 p40 in B6.WT mouse at steady-state and day 2 post-TBI of single cell suspension of isolated LNs stimulated with PMA/ionomycin and Brefeldin A for 4 hr before in culture before flow cytometry measurement on Mφ (CD45^+^CD11c^+^), CD45^+^CD11c^+^CD11b^+^ cells, granulocytes (CD45^+^CD11c^+^CD11b^+^), B cells (CD45^+^CD19^+^) and T cells (CD45^+^CD3ε^+^) and quantification in MFI. **(J)** Mice were administered with IL-12p40 (C17.8) or control rat IgG2a (2A3) antibodies i.p. at 500 µg per dose from day -3 to +2 of allo-HCT. Flow cytometry analysis of donor CD4^+^ T cells (CD90.1^+^CD4^+^) for Ki67 expression from LNs at day 3 of allo-HCT (n = 5-6). Statistical analysis by log-rank test (B and D), two-way ANOVA with Sidak’s multiple comparison test (H and I) and unpaired non-parametric Mann-Whitney test (J), (Mean± SD); \*\**p* < 0.01, \*\*\**p* < 0.001 and \*\*\*\**p* < 0.0001. BECs: blood endothelial cells, TBI: total body irradiation, RNA-seq: RNA sequencing, IFNγ: Interferon gamma, eYFP: enhanced yellow fluorescent protein, Mφ: macrophages, ECs: endothelial cells, MFI; mean fluorescent intensity, PMA: Phorbol-Myristate-Acetate.

To directly test whether SLO ECs prime donor CD4⁺ T cells during aGvHD, we generated mice lacking MHCII expression on Cdh5^+^ endothelial cells (MHCII^ΔCdh5^), including SLO BECs and LECs. At day +3 after allo-HCT, expression of activation markers (CD44, CD25, Ki67) by donor CD4^+^ T cells was comparable in MHCII^ΔCdh5^ recipients and *H2-Ab1*^fl^ littermate control recipients (**Figure S2D-F**). However, survival was strikingly improved (52.1% in MHCII^ΔCdh5^ vs. 20% in littermate controls) (**Figure 3B**), demonstrating an important role for MHCII-expressing ECs in driving aGvHD, even in the presence of intact professional hematopoietic APCs.

Conditional deletion of MHCII selectively in LECs (Prox1-Cre x *H2-Ab1*^fl^ mice), by contrast, provided no survival benefit (**Figure 3C, D**), implicating BECs as the key APC population promoting disease.

BECs expressed high levels of invariant chain (Ii/Cd74), which is required for trafficking of MHCII to MHC class II–containing compartments (MIICs) (**Figure 3E, top**), and the peptide editor H2-M (**Figure 3E, middle**), which facilitates the exchange of CLIP for antigenic peptides (Lazarski et al.), whereas previous studies have shown that these cells express very low levels of H2-O, a negative regulator of H2-M (Rouhani et al.). Staining with an I-A^b^-specific antibody (15G4) confirmed the presence of I-A^b^ molecules bound to Ii degradation intermediates, consistent with active peptide editing (**Figure 3E, bottom**). By contrast, data from earlier studies showed that LECs lacked this machinery (Rouhani et al.).

We then examined how TBI and allo-HCT influence BEC function. Bulk RNA-seq of SLO BECs revealed profound transcriptional remodeling, with principal component analysis (PCA) separating allo-HCT from steady state by 64% variance (**Figure 3F**). Genes involved in MHCII processing, TNF and IFN responses, chemokine signaling, and cytokine–receptor interactions were upregulated after TBI (day 3) and further amplified by allo-HCT (day 3) (**Figure 3G** and **Figure S3A–C**). At the protein level, IFNγ reporter mice demonstrated robust IFNγ induction in myeloid cells and moderate induction in ECs two days post-TBI, whereas B and T cells remained unchanged (**Figure 3H**).

Because interleukin (IL)-12 is a key driver of IFNγ responses (Biron and Tarrio), we profiled IL-12/IL-23p40 expression on LN cells at steady-state or after TBI. LN macrophages and B cells produced abundant IL-12 after TBI (**Figure 3I**), which was associated with activation of JAK–STAT signaling in BECs after TBI and allo-HCT (**Figure S3D**), and increased MHCII expression (**Figure 3G**). Consistent with this pathway, IL-12p40 blockade partially suppressed donor CD4⁺ T cell expansion in LNs during aGvHD initiation (**Figure 3J**). Moreover, systemic IFNγ infusion sufficed to upregulate MHCII on BECs, indicating that autocrine, paracrine, and endocrine IFNγ all converge to regulate endothelial antigen presentation (**Figure S3E**).

Together, these findings show that SLO BECs have the capacity to function as APCs and suggest that BECs sustain MHCII antigen processing and prime donor CD4⁺ T cells, possibly by integrating IFNγ and IL-12 signals, thereby driving alloimmune activation even when professional hematopoietic APCs are intact.

### Activation of allogenic CD4^+^ donor T cells in SLOs can occur independently of hematopoietic antigen presentation

We next asked whether donor CD4⁺ T cell priming in SLOs depends exclusively on hematopoietic APCs, using mice in which MHC class II expression is absent on all CD11c^+^ cells, including classical DCs, CD11c^+^ macrophages, and other CD11c⁺ cells (MHCII^ΔCD11c^) (**Figure S4A**). We compared survival of MHCII^ΔCD11c^ mice after allo-HCT compared with WT littermate controls. In parallel, we used CD11c.DOG mice, in which administration of diphtheria toxin (DT) administration ablates CD11c⁺ cells (**Figure S4B**). At day 3 after allo-HCT, migration of donor CD4⁺ T cells to mesenteric LNs, proliferation, and activation were reduced in MHCII^ΔCD11c^ recipients compared with littermate controls, whereas CD11c-depleted CD11c.DOG mice paradoxically showed increased T cell activation and proliferation (**Figure S4C, D**). Splenic T cells in MHCII^ΔCD11c^ recipient mice, by contrast, still displayed activation by day 3 (**Figure S4D, E**). The hyperactivation of donor T cells in CD11c-depleted CD11c.DOG mice is consistent with a prior report (Koyama et al.). Survival was improved in MHCII^ΔCD11c^ mice, with 80% survival at day 60 post-HCT, compared to 20% survival in *H2-Ab1*^fl^ littermate controls, suggesting that CD11c^+^ APCs are important in the initiation of GvHD after allo-HCT. CD11c.DOG mice developed hyper-acute aGvHD with complete mortality (**Figure S4F**). These findings indicate that although hematopoietic CD11c⁺ APCs enhance alloreactive T cell activation, their absence does not prevent priming and may even unleash pathogenic T cell responses.

To fully ablate MHCII on all hematopoietic APCs, we established MHCII^Δ^ bone marrow (BM) chimeras by transplanting T cell–depleted BM from syngeneic MHCII^Δ^ donors into lethally irradiated WT recipients (MHCII^Δ^ BM ➔ WT) (**Figure 4A, B**). As previously reported (Huang et al.), these chimeras developed a wasting disease within weeks, marked by impaired thymic negative selection and dysregulation of regulatory T (Treg) cells (**Figure 4C, D**), precluding long-term allo-HCT experiments. Nonetheless, in short-term assays, allogeneic CD4⁺ T cells homed to SLOs (spleen, LNs, PPs) and became activated despite the absence of hematopoietic MHCII. However, activation and proliferation were significantly reduced compared to that in WT ➔ WT chimera control recipients (**Figure 4E–4G**). These results demonstrate that residual donor T cell priming occurs in the absence of MHCII expression on hematopoietic APCs, and that non-hematopoietic APCs can provide sufficient antigen presentation to initiate T cell activation during the earliest phase of aGvHD. Based on our earlier results, SLO BECs are likely to be the relevant non-hematopoietic cell population.

**Figure 4:** Alloreactive CD4^+^ T cells activate in the secondary lymphoid organs in the absence of MHC class II on hematopoietic cells in the initiation phase of aGvHD. **(A)** Expression of MHCII on the myeloid APCs (CD45^+^CD11c^+^) of WT BM chimeras versus MHCII^Δ^ BM chimeras at 10 weeks post-syn-HCT. **(B)** Experimental design: TCD BM from the B6.WT or B6.MHCII^Δ^ mice was syn-HCT to myeloablatively irradiated (9 Gy) B6.WT mice, 8-10 weeks post-syn-HCT after successful BM engraftment, mice were again myeloablatively irradiated (9 Gy) and allogenically transplanted with 5x10^6^ TCD BM cells and 5x10^6^ CD4^+^ T cells from FVB and FVB.L2G85 mice, respectively. **(C)** Survival by Kaplan-Meier estimates, absolute weight change and clinical scores of syn-HCT BM chimeras. Data is representative of one experiment (n = 5). **(D)** Spleen size in cm and frequency of Tregs (CD3ε^+^CD4^+^FoxP3^+^) in spleen at day 52 of syn-HCT in BM chimeras. Data is representative of one experiment (n = 4-5). **(E)** Representative e*x vivo* bioluminescent images and quantification on day 3 of allo-HCT. Data is representative of one experiment (n = 3-4). **(F and G)** Flow cytometry analysis of donor T cells (CD90.1^+^CD4^+^) day 3 of allo-HCT. **(F)** CFSE dilution analysis on donor T cells (CD90.1^+^CD4^+^) day 3 of allo-HCT and quantification. Data is representative of one experiment (n = 3-4). **(G)** Expression of CD44 and CD25 in MFI on donor T cells (CD90.1^+^CD4^+^) day 3 of allo-HCT and quantification in MFI fold change in the mLNs. Data is representative of one experiment (n = 3-4). Statistical analysis by log-rank test (C), unpaired parametric, 2-tailed Student’s t test (D, F, G) and two-way ANOVA with Sidak’s multiple comparison test (E) (Mean± SD); \**p* < 0.05, \*\**p* < 0.01, \*\*\**p* < 0.001 and \*\*\*\**p* < 0.0001. APCs: antigen presenting cells, syn-HCT: syngeneic HCT, allo-HCT: allogeneic HCT, MFI: mean fluorescent intensity.

### SLO BECs function as non-hematopoietic APCs that activate allogeneic CD4^+^ T cells

Because MHCII^Δ^ ➔ WT BM chimeras were unsuitable to study endothelial antigen presentation, we turned to a genetic strategy that selectively preserves MHCII on SLO ECs. Revisiting our scRNA-seq data, we confirmed that *Vav1* is expressed in all hematopoietic cells but is absent in ECs (Ogilvy et al.) (**Figure 5A**). We therefore generated mice that lacked MHCII on hematopoietic lineages but retained expression on ECs (MHCII^ΔVav1^), enabling us to directly test EC–mediated priming (**Figure S5A**). LN BECs sorted from MHCII^ΔVav1^ mice efficiently processed DQ-OVA *in vitro*, demonstrating intact antigen processing capacity (**Figure 5B**), and upregulated MHC class II in response to IFNγ infusion *in vivo* (**Figure 5C**). MHCII^ΔVav1^ recipient mice were only modestly protected in the allo-HCT model, with 33.3% survival at day 60 compared to 0% in littermate *H2-Ab1*^fl^ control recipients (**Figure 5D**), indicating that MHC class II-expressing ECs are sufficient to drive GvHD after allo-HCT.

**Figure 5:** Allogeneic CD4^+^ T cells activate by the non-hematopoietic cells of LNs. **(A)** Expression of *Vav1* across LN ECs evaluated by scRNA-seq on UMAP plot. **(B)** DQ-OVA incubated with FACS sorted blood endothelial cells from B6.MHCII^ΔVav1^ mice and cultured for 3 h at 4°C (blue) or 37°C (red) in histogram, followed by analysis of processed DQ-OVA under the blue laser (488 nm), BL-1 and quantification of DQ-OVA processing of blood endothelial cells from C57BL/6 (blue) vs B6.MHCII^ΔVav1^ (red) in geometrical MFI fold change. **(C)** Expression of MHCII on LN blood endothelial cells post IFNγ infusion in B6.MHCII^ΔVav1^ mice. Mice were s.c. injected with IFNγ (3 µg) or DPBS for 3 days followed by MHCII expression analysis on blood endothelial cells (CD45^-^CD24^-^gp38^-^CD31^+^) on flow cytometer and quantification in fold-change MFI. **(D)** Myeloablatively irradiated (9 Gy) B6.*H2-Ab1*^fl^ and B6.MHCII^ΔVav1^ mice were i.v. transplanted with 5x10^6^ TCD BM and 6x10^5^ CD4^+^ T cells from FVB. Survival by Kaplan-Meier estimates, clinical scores combined from 3 replicate experiments. **(E and F)** MLR of BALB/c CD4^+^ T cells (H-2K^d+^CD4^+^) co-cultured with irradiated non-hematopoietic lymph node stromal cells (LNSCs) from (BALB/c, C57BL/6, B6.MHCII^Δ^ and B6.MHCII^ΔVav1^ mice), magnetically depleted of CD45^+^ cells, analyzed at day 5 of co-culture. **(E)** Flow cytometry expression analysis of CD44, CD25 and CXCR3 shown in histogram quantification in MFI fold change. Data are representative of 2 replicate experiments (n = 3-4). **(F)** Ki67^+^ expression on BALB/c CD4^+^ T cells (H-2K^d+^CD4^+^) in MLR and quantification. Data are representative of 2 replicate experiments (n = 3-4). **(G)** Experimental design: mLNs from B6.MHCII^Δ^ recipients were surgically removed and transplanted with mLNs from B6.MHCII^ΔVav1^ mouse and mLNs from B6.MHCII^Δ^ into B6.MHCII^Δ^ recipients as controls, 6-8 weeks after surgery MHC major mismatch aGvHD was induced by myeloablative irradiation (9 Gy) and i.v. injection of 5x10^6^ TCD BM cells and 5x10^6^ CD4^+^ T cells from FVB mouse. **(H)** Expression of TNFα and IFNγ on FVB donor CD4^+^ T cells (CD90.1^+^CD4^+^) from mLNs of B6.MHCII^ΔVav1^ (positive control), B6.MHCII^Δ^ mouse with B6.MHCII^Δ^ mLNs and B6.MHCII^Δ^ mouse with B6.MHCII^ΔVav1^ mLNs at day 12 of allo-HCT. Single cell suspension of isolated mLNs was stimulated with PMA/ionomycin with Brefeldin A for 4 hr before in culture before flow cytometry measurement. Data are representative of 2 replicate experiments (n = 3). Statistical analysis by unpaired parametric, 2-tailed Student’s t test (B and C), log-rank test (D), Ordinary one-way ANOVA, adjusted for multiple comparisons with Tukey’s multiple-comparison test (E, F and H), (Mean± SD); \**p* < 0.05 and \*\**p* < 0.01. DQ-OVA: BODIPY-conjugated OVA, FACS: Fluorescence activated cell sorting, TCD: T cell-depleted, MLR: Mixed lymphocyte reaction, TNFα: Tumor necrosis factor alpha, IFNγ: Interferon gamma, PMA: Phorbol-Myristate-Acetate.

To test the ability of BECs to directly prime allogeneic CD4⁺ T cells, we did polyclonal mixed lymphocyte reactions with BALB/c CD4⁺ T cells and CD45⁺-depleted LN stromal cells from B6.MHCII^ΔVav1^ mice, B6.MHCII^Δ^, or WT B6 control mice. Expression of CD44, CD25, and CXCR3 on CD4⁺ T cells was comparable in cells cultured with stromal cells from B6.MHCII^ΔVav1^ or WT B6 mice, whereas expression was significantly lower in CD4⁺ T cells cultured with B6.MHCII^Δ^ stromal cells (**Figure 5E**). Effector-to-naïve cell ratios and proliferation of CD4⁺ T cells showed similar differences (**Figure 5F** and **Figure S5B**). These data show that stromal cells, including BEC, can drive allogeneic CD4⁺ T cell activation in the absence of hematopoietic APCs.

To further assess *in vivo* T cell priming by ECs, we transplanted mesenteric LNs from B6.MHCII^ΔVav1^ or MHCII^Δ^ donors into MHCII^Δ^ recipients and performed allo-HCT 6-8 weeks later (**Figure 5G**). On day 12 after allo-HCT, a higher number of LN CD4⁺ T cells from mice engrafted with B6.MHCII^ΔVav1^ LNs (or intact B6.MHCII^ΔVav1^ mice) co-expressed TNFα and IFNγ compared to mice engrafted with MHCII^Δ^ LNs (**Figure 5H**). We verified these data using an independent model, in which OVA-specific T cells (OT-II) were transferred into irradiated recipient WT mice (no antigen) or recipient mice expressing membrane-bound OVA (B6.Act-mOVA mice or B6 mice engrafted with mesenteric LNs from Act-mOVA donors). OT-II cells underwent robust activation, proliferation, and effector differentiation in OVA-expressing recipient mice, but not in WT B6 recipients (**Figure S6A–S6E**).

Together, these findings establish that SLO BECs, through their intrinsic MHCII expression and antigen-processing machinery, are sufficient to prime and activate alloreactive CD4⁺ T cells *in vivo*, even in the absence of hematopoietic APCs.

### SLO BECs provide sufficient MHCII-dependent antigen presentation to drive lethal aGvHD

To establish the clinical relevance of endothelial antigen presentation in aGvHD, we generated MHCII^ΔVav1ΔCdh5^ mice, which lack MHC class II on both hematopoietic and endothelial cells and used these mice as recipients of allo-HCT, with BALB/c Nur77-eGFP reporter mice as donors (Moran et al.). We found that donor CD4⁺ T cells mounted strong T cell receptor (TCR) signaling in MHCII^ΔVav1^ recipients, whereas signaling was abolished in MHCII^ΔVav1ΔCdh5^ mice, mirroring the absence of responses in MHCII^Δ^ controls (**Figure 6A**). Using an *in vitro* activation T cell activation assay (IL-2 production by BO-97.10 T cell hybridomas; (Hugo et al.), co-culture of hybridoma cells with LN stromal cells from MHCII^ΔVav1^ mice triggered significantly higher IL-2 production than did co-culture with stromal cells from MHCII^ΔVav1ΔCdh5^ mice, although reduced compared to stromal cells from MHCII-competent littermates (**Figure S5C**). Thus, SLO ECs present antigen in an MHCII-dependent manner.

**Figure 6:** LN blood endothelial cells initiates TCR signalling in allogeneic CD4^+^ T cells and exacerbates aGvHD. **(A)** B6.*H2-Ab1*^fl^, B6.MHCII^Δ^, B6.MHCII^ΔVav1^ and B6.MHCII^ΔVav1ΔCdh5^ recipient mice were myeloablatively irradiated with 9 Gy and i.v. transplanted with 5x10^6^ TCD BM and enriched splenic 2x10^6^ CD4^+^ T cells from BALB/c and C.Nur77-eGFP mice, respectively. Expression of Nur77-eGFP on donor CD4^+^ T cells (H-2K^d+^CD4^+^) from mLNs of B6.*H2-Ab1*^fl^ (positive control), B6.MHCII^Δ^ (negative control), B6.MHCII^ΔVav1^ and B6.MHCII^ΔVav1^ ^Cdh5^ recipient mice at day 3.5 of allo-HCT and quantification in fold-change MFI. Data are representative of 2 replicate experiments (n = 4-5). **(B)** Myeloablatively irradiated (9 Gy) B6.*H2-Ab1*^fl^, B6.MHCII^ΔCdh5^, B6.MHCII^ΔVav1^ and B6.MHCII^ΔVav1ΔCdh5^ recipient mice were i.v. transplanted with 5x10^6^ TCD BM and enriched splenic 1.2x10^6^ CD4^+^ T cells from FVB. Survival by Kaplan-Meier estimates and clinical scores combined from 3 replicate experiments. **(C)** Bulk-RNA seq of FACS sorted CD90.1^+^CD4^+^ T cells from B6.MHCII^ΔVav1^ and B6.MHCII^ΔVav1Cdh5^ at day 22 of allo-HCT. Heatmap of significant differentially expressed genes (DEGs, log_2_ fold change ≥ 0.75, *q* ≤ 0.05) relating to T cell activation and suppression pathways on allogeneic CD4^+^ T cells from spleen and LNs. **(D)** RT^2^ Profiler PCR array of CD4^+^ T cells from B6.MHCII^ΔVav1^ and B6.MHCII^ΔVav1^ ^Cdh5^ at day 22 of allo-HCT. Total RNA was isolated from CD4^+^ T cells enriched from secondary lymphoid organs and screened by RT Profiler array of 84 T-/B-cell activation genes by real-time PCR. The array data was normalized with housekeeping gene panel *Actb, B2m, Gapdh, Gusb* and *Hsp90ab1*. Differential gene expression represented as a scatter plot comparing the normalized expression of every gene on the array between the two select groups by plotting them against one another to visualize gene-expression changes. The central line indicates unchanged gene expression. The blue dotted lines indicate the selected fold-regulation threshold. Data points beyond the dotted lines in the upper left and lower right sections meet the selected fold-regulation threshold. Data are representative of one experiment (n = 3-4). **(E)** Myeloablatively irradiated (9 Gy) B6.*H2-Ab1*^fl^, B6.MHCII^ΔVav1ΔVil^ and B6.MHCII^ΔVav1ΔVilΔCdh5^ recipient mice were i.v. transplanted with 5x10^6^ TCD BM and enriched splenic 1.2x10^6^ CD4^+^ T cells from FVB. Survival by Kaplan-Meier estimates and clinical scores combined from 2 replicate experiments. Statistical analysis by One-way ANOVA, adjusted for multiple comparisons with Tukey’s multiple-comparison test (A), log-rank test (B, F), Two-way ANOVA, adjusted for multiple comparisons with Sidak’s multiple-comparison test (C), (Mean± SD); \**p* < 0.05, \*\**p* < 0.01 and \*\*\**p* < 0.001. TCR: T cell receptor, eGFP: Enhanced green fluorescent protein, MFI: Mean fluorescent intensity.

Because aGvHD targets visceral tissues, we next tested whether ECs from liver or lung could act similarly. Unlike SLO BECs, liver and lung ECs from MHCII^ΔVav1^ mice failed to activate or expand CD4⁺ T cells (**Figure S7A–F**), indicating that SLO BECs uniquely function as APCs.

After allo-HCT, MHCII^ΔVav1ΔCdh5^ recipient mice developed only mild disease and showed significantly improved survival at day 60 compared to MHCII^ΔVav1^ recipients, in which class II expression was restricted to ECs (79% survival vs 44%) or to *H2-Ab1*^fl^ littermate controls (10% survival). Survival of MHCII^ΔCdh5^ recipient mice, in which MHC class II expression was restricted to hematopoietic cells, was similar to that of MHCII^ΔVav1^ recipients (33% vs 44%). (**Figure 6B**). RNA-seq of donor CD4⁺ T cells at day 22 after allo-HCT revealed transcriptional reprogramming in MHCII^ΔVav1ΔCdh5^ recipients, with downregulation of TCR signaling genes (*Nr4a1/2/3, Tbx21*), effector molecules (*Gzmb, Fas, Ifng, Tnf*), and interferon pathway regulators (*Irf1, Irf4*), accompanied by modest upregulation of anergy-associated transcripts (*Cd160, Cd244a, Klrg1*) as compared to T cells from MHCII^ΔVav1^ recipients (**Figure 6C**). qPCR confirmed reduced expression of BM homing and proinflammatory genes (*Cxcr4, H60a, Il6*) alongside enrichment for markers of naïve or undifferentiated states (*Nos2, Cr2, Cd79a, Cd38*) (**Figure 6D**) (DeRogatis et al.; Niedbala et al.; Pekalski et al.).

Finally, to assess a potential contribution by intestinal epithelial cells (IEC), as had been shown in a prior study (Koyama et al.), we compared B6.MHCII^ΔVav1ΔVil^ mice (lacking MHCII on hematopoietic and IEC cells) with B6.MHCII^ΔVav1ΔVilΔCdh5^ mice (additionally lacking MHCII on ECs). Strikingly, B6.MHCII^ΔVav1ΔVilΔCdh5^ recipient mice were strongly protected (83% survival at day 60) compared to B6.MHCII^ΔVav1ΔVil^ mice (36%) or littermate controls (7%; **Figure 6E**). These results demonstrate that MHCII-dependent antigen presentation by SLO ECs is both necessary and sufficient to drive lethal allogeneic CD4⁺ T cell responses, independent of hematopoietic APCs or IECs.

## Discussion

Donor T cell priming and activation are pivotal in determining the outcome of allo-HCT, whether toward graft-versus-leukemia or GvHD. While professional hematopoietic APCs have long been recognized as key drivers of donor T cell activation in this setting, our study identifies a novel role for SLO BECs in MHC class II-dependent antigen presentation during the initiation phase of aGvHD. First, we confirmed the importance of SLOs in allogeneic T cell priming by showing that donor CD4⁺ T cells are initially confined to SLOs (Beilhack et al.; Brede et al.), whereas they appear in peripheral target tissues, such as the intestinal lamina propria, only at later timepoints after having undergone activation and proliferation.

Although professional APCs, particularly DCs, are central to GvHD initiation (Duffner et al.), prior studies have shown that eliminating MHC class II–dependent antigen presentation by hematopoietic cells does not fully prevent aGvHD (Koyama et al.; Li et al.; Toubai et al.), suggesting a contribution of non-hematopoietic APCs in allo-T cell priming. Recent work has shown that presentation of commensal gut bacterial by intestinal epithelial cells (IECs) exacerbates gut pathology during the effector phase of aGvHD (Koyama et al.). However, these cells are unlikely to contribute to the initiation phase of allogeneic T cell priming given that naïve T cells must transit through SLOs (which are devoid of IECs) in order to become activated and express gut-homing receptors (Beilhack et al.; Johansson-Lindbom et al.; Mora et al.), and interfering with T cell access to SLOs, either by blocking entry or depleting SLOs entirely, prevents aGvHD (Beilhack et al.; Coghill et al.). Together, the evidence points to an important role for non-hematopoietic APCs within SLOs in initiating donor T cell priming during aGvHD, which remains intact in the absence of functional hematopoietic APCs.

Other non-hematopoietic stromal cells in SLOs possess APC function, including fibroblastic reticular cells (FRCs) and LECs, but we felt that these populations were unlikely to be relevant in our model. First, many studies have shown that these populations primarily function to enforce tolerance via secretion of immunoregulatory molecules such as prostaglandin E2 and nitric oxide, expression of inhibitory receptors such as PD-L1, and activation of Treg cells (Baptista et al.; Dubrot et al.; Dubrot et al.; Gkountidi et al.; Krishnamurty and Turley; Lukacs-Kornek et al.; Rouhani et al.; Schaeuble et al.; Shaikh et al.; Tewalt et al.). In addition, although LECs express MHCII and invariant chain, they lack the capacity to present antigen via MHC class II, likely due to lack of H2-M expression and thus an inability to load antigenic peptides onto MHCII molecules (Rouhani et al.). Finally, prior work from our group showed that selective deletion of MHC class II in FRCs exacerbated aGvHD due in part to defective Treg cell maintenance (Shaikh et al.). As such, in this work we focused our attention on the other primary non-hematopoietic cell type that resides in SLOs: BECs.

In our *in vivo* allo-HCT model, mice lacking MHC class II only in the EC compartment (MHCII^ΔCdh5^), but not those lacking MHCII only in LECs (MHCII^ΔProx1^), were partially protected against lethal aGvHD, suggesting a contributory role for BECs in initiating pathogenic allo-T cells responses. Consistent with this, our *ex vivo* data confirm and extend prior studies showing that BECs possess the machinery for antigen presentation, including high levels of invariant chain (Ii/CD74) and the peptide editor H2-M, as well as co-stimulatory molecules, endowing them with the capacity to prime naïve CD4⁺ T cells and drive memory responses (Abrahimi et al.; Rothermel et al.). By contrast, BECs from peripheral organs such as liver and lung failed to prime naïve T cells despite expressing MHC class II, likely due to the absence of the co-stimulatory molecules CD80 and CD86, underscoring tissue-specific functional heterogeneity (Amersfoort et al.; Carambia et al.; Kalucka et al.; Kreisel et al.).

Our data also corroborated an important role for hematopoietic cells in the initiation of aGvHD, given that survival was comparably increased in recipient mice lacking MHCII in hematopoietic cells (MHCII^ΔVav1^) as in those lacking MHCII in ECs (de Boer et al.; Rodriguez et al). However, the fact that aGvHD still developed in mice lacking hematopoietic MHCII expression indicates an important role for MHCII-expressing ECs. Indeed, combined deletion of MHCII in both hematopoietic and endothelial cells (MHCII^ΔVav1ΔCdh5^) mice drastically reduced GvHD severity, improved survival. In totality, the data suggest that both hematopoietic APCs and BECs are involved in the initiation of pathogenic T cell responses after allo-HCT, and further studies are needed to disentangle their relative contributions in the context of a fully intact immune system.

Mechanistically, we provide evidence that local production of IFNγ promotes antigen presentation by BECs. IFNγ was produced by multiple cells types, including endothelial cells, in response to pre-transplant myeloablative conditioning (TBI) in our model, and infusion of IFNγ upregulated expression of MHCII on ECs *in vivo*, consistent with prior findings (Pober et al.). Clinically, systemic IFNγ blockade in humans is limited by toxicity (Burman et al.; Mauermann et al.), whereas targeting IL-12, an upstream driver of IFNγ, is feasible and has been shown to be protective in mouse models of GvHD (Feagan et al.; Pidala et al.). Indeed, blockade of IL-12 with the FDA-approved anti-IL-12/23 p40 antibody Ustekinumab has shown benefit in patients undergoing allo-HCT in one trial (Pidala et al.), with other trials currently underway (e.g., NCT04572815). Definitive evidence of a pivotal role for IL-12 and IFNγ in our model will require further studies, for example using in mice lacking the IFNγ receptor in ECs.

Overall, our findings reveal a previously unrecognized role of SLO BECs as non-hematopoietic APCs that contribute to allogeneic CD4⁺ T cell priming and the initiation of lethal GvHD, with the IL-12/IFNγ axis implicated as a key regulator of MHC class II expression on BECs. These findings not only reshape the paradigm of alloimmune priming but also corroborate clinically actionable targets to prevent or mitigate GvHD at the earliest stage. Beyond transplantation, these results may have broad implications for other inflammatory and autoimmune conditions initiated in lymphoid organs.

## Supporting information

Main figures

## ACKNOWLEDGEMENTS

This work was supported by grants from the German research council (DFG) to HS (SFB221, 324392634; 549527997), AB (SFB221, 324392634; GRK2157 P1, 270563345), Bayerische Forschungsstifung (Fortither, WP2TP3), and the Europäische Fonds für Regionale Entwicklung (EFRE; Center for Personalized Molecular Immunotherapy). scRNA-Seq and bulk RNA-seq experiments were performed at the Core Unit Systemmedizin, University of Würzburg. We would like to express our gratitude to Caroline Graf and Estibaliz Arellano Viera for excellent technical support.

## AUTHOR CONTRIBUTIONS

HS designed, planned, and carried out the experiments and analyzed and interpreted the data. MH, JP, ZM, and AR analyzed sequencing data. JGV, MAGK, KJ, HY and JPJM assisted with the experiments. PW and ZA performed and analyzed 3D-LSFM and confocal data, respectively. JP planned and performed the scRNA-seq experiments. MBH, HE, AZ and JH provided intellectual support and assisted in editing the manuscript. HS and AB designed the study, interpreted the data, and wrote the manuscript. All authors read and discussed the manuscript.

## REFERENCES

Abrahimi, P., Qin, L., Chang, W.G., Bothwell, A.L., Tellides, G., Saltzman, W.M., and Pober, J.S. (2016). Blocking MHC class II on human endothelium mitigates acute rejection. JCI Insight 1.

Amersfoort, J., Eelen, G., and Carmeliet, P. (2022). Immunomodulation by endothelial cells - partnering up with the immune system? Nat Rev Immunol 22, 576–588.

Bajenoff, M., Egen, J.G., Koo, L.Y., Laugier, J.P., Brau, F., Glaichenhaus, N., and Germain, R.N. (2006). Stromal cell networks regulate lymphocyte entry, migration, and territoriality in lymph nodes. Immunity 25, 989–1001.

Baptista, A.P., Roozendaal, R., Reijmers, R.M., Koning, J.J., Unger, W.W., Greuter, M., Keuning, E.D., Molenaar, R., Goverse, G., Sneeboer, M.M., et al. (2014). Lymph node stromal cells constrain immunity via MHC class II self-antigen presentation. Elife 3.

Bauerlein, C.A., Riedel, S.S., Baker, J., Brede, C., Garrote, A.L., Chopra, M., Ritz, M., Beilhack, G.F., Schulz, S., Zeiser, R., et al. (2013). A diagnostic window for the treatment of acute graft-versus-host disease prior to visible clinical symptoms in a murine model. BMC Med 11, 134.

Beilhack, A., Schulz, S., Baker, J., Beilhack, G.F., Nishimura, R., Baker, E.M., Landan, G., Herman, E.I., Butcher, E.C., Contag, C.H., and Negrin, R.S. (2008). Prevention of acute graft-versus-host disease by blocking T-cell entry to secondary lymphoid organs. Blood 111, 2919–2928.

Beilhack, A., Schulz, S., Baker, J., Beilhack, G.F., Wieland, C.B., Herman, E.I., Baker, E.M., Cao, Y.A., Contag, C.H., and Negrin, R.S. (2005). In vivo analyses of early events in acute graft-versus-host disease reveal sequential infiltration of T-cell subsets. Blood 106, 1113–1122.

Biron, C.A., and Tarrio, M.L. (2015). Immunoregulatory cytokine networks: 60 years of learning from murine cytomegalovirus. Med Microbiol Immunol 204, 345–354.

Brede, C., Friedrich, M., Jordan-Garrote, A.L., Riedel, S.S., Bauerlein, C.A., Heinze, K.G., Bopp, T., Schulz, S., Mottok, A., Kiesel, C., et al. (2012). Mapping immune processes in intact tissues at cellular resolution. J Clin Invest 122, 4439–4446.

Burman, A.C., Banovic, T., Kuns, R.D., Clouston, A.D., Stanley, A.C., Morris, E.S., Rowe, V., Bofinger, H., Skoczylas, R., Raffelt, N., et al. (2007). IFNgamma differentially controls the development of idiopathic pneumonia syndrome and GVHD of the gastrointestinal tract. Blood 110, 1064–1072.

Carambia, A., Freund, B., Schwinge, D., Heine, M., Laschtowitz, A., Huber, S., Wraith, D.C., Korn, T., Schramm, C., Lohse, A.W., et al. (2014). TGF-beta-dependent induction of CD4(+)CD25(+)Foxp3(+) Tregs by liver sinusoidal endothelial cells. J Hepatol 61, 594–599.

Card, C.M., Yu, S.S., and Swartz, M.A. (2014). Emerging roles of lymphatic endothelium in regulating adaptive immunity. J Clin Invest 124, 943–952.

Chen, L., Li, J., Wang, F., Dai, C., Wu, F., Liu, X., Li, T., Glauben, R., Zhang, Y., Nie, G., et al. (2016). Tie2 Expression on Macrophages Is Required for Blood Vessel Reconstruction and Tumor Relapse after Chemotherapy. Cancer Res 76, 6828–6838.

Coghill, J.M., Sarantopoulos, S., Moran, T.P., Murphy, W.J., Blazar, B.R., and Serody, J.S. (2011). Effector CD4+ T cells, the cytokines they generate, and GVHD: something old and something new. Blood 117, 3268–3276.

Cohen, J.N., Tewalt, E.F., Rouhani, S.J., Buonomo, E.L., Bruce, A.N., Xu, X., Bekiranov, S., Fu, Y.X., and Engelhard, V.H. (2014). Tolerogenic properties of lymphatic endothelial cells are controlled by the lymph node microenvironment. PLoS One 9, e87740.

DeRogatis, J.M., Neubert, E.N., Viramontes, K.M., Henriquez, M.L., Nicholas, D.A., and Tinoco, R. (2023). Cell-Intrinsic CD38 Expression Sustains Exhausted CD8(+) T Cells by Regulating Their Survival and Metabolism during Chronic Viral Infection. J Virol 97, e0022523.

Dubrot, J., Duraes, F.V., Harle, G., Schlaeppi, A., Brighouse, D., Madelon, N., Gopfert, C., Stokar-Regenscheit, N., Acha-Orbea, H., Reith, W., et al. (2018). Absence of MHC-II expression by lymph node stromal cells results in autoimmunity. Life Sci Alliance 1, e201800164.

Dubrot, J., Duraes, F.V., Potin, L., Capotosti, F., Brighouse, D., Suter, T., LeibundGut-Landmann, S., Garbi, N., Reith, W., Swartz, M.A., and Hugues, S. (2014). Lymph node stromal cells acquire peptide-MHCII complexes from dendritic cells and induce antigen-specific CD4(+) T cell tolerance. J Exp Med 211, 1153–1166.

Duffner, U.A., Maeda, Y., Cooke, K.R., Reddy, P., Ordemann, R., Liu, C., Ferrara, J.L., and Teshima, T. (2004). Host dendritic cells alone are sufficient to initiate acute graft-versus-host disease. J Immunol 172, 7393–7398.

Feagan, B.G., Sandborn, W.J., Gasink, C., Jacobstein, D., Lang, Y., Friedman, J.R., Blank, M.A., Johanns, J., Gao, L.L., Miao, Y., et al. (2016). Ustekinumab as Induction and Maintenance Therapy for Crohn’s Disease. N Engl J Med 375, 1946–1960.

Flamann, C.S., Shaikh, H., Matos, C., Kreutz, M., Ali, H., Kern, M.A., Buttner-Herold, M., Jacobs, B., Volkl, S., Lischer, C., et al. (2025). Augmented CD47 expression impairs alloreactive T-cell clearance after allo-HCT. Blood.

Gkountidi, A.O., Garnier, L., Dubrot, J., Angelillo, J., Harle, G., Brighouse, D., Wrobel, L.J., Pick, R., Scheiermann, C., Swartz, M.A., and Hugues, S. (2021). MHC Class II Antigen Presentation by Lymphatic Endothelial Cells in Tumors Promotes Intratumoral Regulatory T cell-Suppressive Functions. Cancer Immunol Res 9, 748–764.

Hashimoto, D., Chow, A., Greter, M., Saenger, Y., Kwan, W.H., Leboeuf, M., Ginhoux, F., Ochando, J.C., Kunisaki, Y., van Rooijen, N., et al. (2011). Pretransplant CSF-1 therapy expands recipient macrophages and ameliorates GVHD after allogeneic hematopoietic cell transplantation. J Exp Med 208, 1069–1082.

Huang, W., Qi, Q., Hu, J., Huang, F., Laufer, T.M., and August, A. (2014). Dendritic cell-MHC class II and Itk regulate functional development of regulatory innate memory CD4+ T cells in bone marrow transplantation. J Immunol 192, 3435–3441.

Hugo, P., Kappler, J.W., Godfrey, D.I., and Marrack, P.C. (1992). A cell line that can induce thymocyte positive selection. Nature 360, 679–682.

Hulsdunker, J., Ottmuller, K.J., Neeff, H.P., Koyama, M., Gao, Z., Thomas, O.S., Follo, M., Al-Ahmad, A., Prinz, G., Duquesne, S., et al. (2018). Neutrophils provide cellular communication between ileum and mesenteric lymph nodes at graft-versus-host disease onset. Blood 131, 1858–1869.

Jalkanen, S., and Salmi, M. (2020). Lymphatic endothelial cells of the lymph node. Nat Rev Immunol 20, 566–578.

Johansson-Lindbom, B., Svensson, M., Pabst, O., Palmqvist, C., Marquez, G., Forster, R., and Agace, W.W. (2005). Functional specialization of gut CD103+ dendritic cells in the regulation of tissue-selective T cell homing. J Exp Med 202, 1063–1073.

Junt, T., Scandella, E., and Ludewig, B. (2008). Form follows function: lymphoid tissue microarchitecture in antimicrobial immune defence. Nat Rev Immunol 8, 764–775.

Kalucka, J., de Rooij, L., Goveia, J., Rohlenova, K., Dumas, S.J., Meta, E., Conchinha, N.V., Taverna, F., Teuwen, L.A., Veys, K., et al. (2020). Single-Cell Transcriptome Atlas of Murine Endothelial Cells. Cell 180, 764–779 e720.

Kambayashi, T., and Laufer, T.M. (2014). Atypical MHC class II-expressing antigen-presenting cells: can anything replace a dendritic cell? Nat Rev Immunol 14, 719–730.

Koyama, M., and Hill, G.R. (2016). Alloantigen presentation and graft-versus-host disease: fuel for the fire. Blood 127, 2963–2970.

Koyama, M., and Hill, G.R. (2019). The primacy of gastrointestinal tract antigen-presenting cells in lethal graft-versus-host disease. Blood 134, 2139–2148.

Koyama, M., Hippe, D.S., Srinivasan, S., Proll, S.C., Miltiadous, O., Li, N., Zhang, P., Ensbey, K.S., Hoffman, N.G., Schmidt, C.R., et al. (2023). Intestinal microbiota controls graft-versus-host disease independent of donor-host genetic disparity. Immunity 56, 1876–1893 e1878.

Koyama, M., Kuns, R.D., Olver, S.D., Raffelt, N.C., Wilson, Y.A., Don, A.L., Lineburg, K.E., Cheong, M., Robb, R.J., Markey, K.A., et al. (2011). Recipient nonhematopoietic antigen-presenting cells are sufficient to induce lethal acute graft-versus-host disease. Nat Med 18, 135–142.

Koyama, M., Mukhopadhyay, P., Schuster, I.S., Henden, A.S., Hulsdunker, J., Varelias, A., Vetizou, M., Kuns, R.D., Robb, R.J., Zhang, P., et al. (2019). MHC Class II Antigen Presentation by the Intestinal Epithelium Initiates Graft-versus-Host Disease and Is Influenced by the Microbiota. Immunity 51, 885–898 e887.

Kreisel, D., Richardson, S.B., Li, W., Lin, X., Kornfeld, C.G., Sugimoto, S., Hsieh, C.S., Gelman, A.E., and Krupnick, A.S. (2010). Cutting edge: MHC class II expression by pulmonary nonhematopoietic cells plays a critical role in controlling local inflammatory responses. J Immunol 185, 3809–3813.

Krishnamurty, A.T., and Turley, S.J. (2020). Lymph node stromal cells: cartographers of the immune system. Nat Immunol 21, 369–380.

Lazarski, C.A., Chaves, F.A., and Sant, A.J. (2006). The impact of DM on MHC class II-restricted antigen presentation can be altered by manipulation of MHC-peptide kinetic stability. J Exp Med 203, 1319–1328.

Le, D.D., Jordán Garrote, A.-L., Maria, R., Ottmüller, K.J., Shaikh, H., Qureischi, M., Scheller, L., Steinfatt, T., Brandl, A., Hartweg, J., et al. (2018). Tissue Derived Non-Classical Monocyte Derived Host Macrophages Protect Against Murine Intestinal Acute Graft-Versus-Host Disease. Blood 132, 3315–3315.

Li, H., Demetris, A.J., McNiff, J., Matte-Martone, C., Tan, H.S., Rothstein, D.M., Lakkis, F.G., and Shlomchik, W.D. (2012). Profound depletion of host conventional dendritic cells, plasmacytoid dendritic cells, and B cells does not prevent graft-versus-host disease induction. J Immunol 188, 3804–3811.

Londei, M., Bottazzo, G.F., and Feldmann, M. (1985). Human T-cell clones from autoimmune thyroid glands: specific recognition of autologous thyroid cells. Science 228, 85–89.

Lukacs-Kornek, V., Malhotra, D., Fletcher, A.L., Acton, S.E., Elpek, K.G., Tayalia, P., Collier, A.R., and Turley, S.J. (2011). Regulated release of nitric oxide by nonhematopoietic stroma controls expansion of the activated T cell pool in lymph nodes. Nat Immunol 12, 1096–1104.

Mauermann, N., Burian, J., von Garnier, C., Dirnhofer, S., Germano, D., Schuett, C., Tamm, M., Bingisser, R., Eriksson, U., and Hunziker, L. (2008). Interferon-gamma regulates idiopathic pneumonia syndrome, a Th17+CD4+ T-cell-mediated graft-versus-host disease. Am J Respir Crit Care Med 178, 379–388.

Mora, J.R., Bono, M.R., Manjunath, N., Weninger, W., Cavanagh, L.L., Rosemblatt, M., and Von Andrian, U.H. (2003). Selective imprinting of gut-homing T cells by Peyer’s patch dendritic cells. Nature 424, 88–93.

Moran, A.E., Holzapfel, K.L., Xing, Y., Cunningham, N.R., Maltzman, J.S., Punt, J., and Hogquist, K.A. (2011). T cell receptor signal strength in Treg and iNKT cell development demonstrated by a novel fluorescent reporter mouse. J Exp Med 208, 1279–1289.

Mueller, J.P.J., Dobosz, M., O’Brien, N., Abdoush, N., Giusti, A.M., Lechmann, M., Osl, F., Wolf, A.K., Arellano-Viera, E., Shaikh, H., et al. (2023). ROCKETS - a novel one-for-all toolbox for light sheet microscopy in drug discovery. Front Immunol 14, 1034032.

Muhlethaler-Mottet, A., Otten, L.A., Steimle, V., and Mach, B. (1997). Expression of MHC class II molecules in different cellular and functional compartments is controlled by differential usage of multiple promoters of the transactivator CIITA. EMBO J 16, 2851–2860.

Niedbala, W., Cai, B., and Liew, F.Y. (2006). Role of nitric oxide in the regulation of T cell functions. Ann Rheum Dis 65 Suppl 3, iii37–40.

Ogilvy, S., Metcalf, D., Gibson, L., Bath, M.L., Harris, A.W., and Adams, J.M. (1999). Promoter elements of vav drive transgene expression in vivo throughout the hematopoietic compartment. Blood 94, 1855–1863.

Onder, L., Morbe, U., Pikor, N., Novkovic, M., Cheng, H.W., Hehlgans, T., Pfeffer, K., Becher, B., Waisman, A., Rulicke, T., et al. (2017). Lymphatic Endothelial Cells Control Initiation of Lymph Node Organogenesis. Immunity 47, 80–92 e84.

Pekalski, M.L., Garcia, A.R., Ferreira, R.C., Rainbow, D.B., Smyth, D.J., Mashar, M., Brady, J., Savinykh, N., Dopico, X.C., Mahmood, S., et al. (2017). Neonatal and adult recent thymic emigrants produce IL-8 and express complement receptors CR1 and CR2. JCI Insight 2.

Perez-Shibayama, C., Gil-Cruz, C., and Ludewig, B. (2019). Fibroblastic reticular cells at the nexus of innate and adaptive immune responses. Immunol Rev 289, 31–41.

Pezoldt, J., Pasztoi, M., Zou, M., Wiechers, C., Beckstette, M., Thierry, G.R., Vafadarnejad, E., Floess, S., Arampatzi, P., Buettner, M., et al. (2018). Neonatally imprinted stromal cell subsets induce tolerogenic dendritic cells in mesenteric lymph nodes. Nat Commun 9, 3903.

Pidala, J., Beato, F., Kim, J., Betts, B., Jim, H., Sagatys, E., Levine, J.E., Ferrara, J.L.M., Ozbek, U., Ayala, E., et al. (2018). In vivo IL-12/IL-23p40 neutralization blocks Th1/Th17 response after allogeneic hematopoietic cell transplantation. Haematologica 103, 531–539.

Pober, J.S., Gimbrone, M.A., Jr., Cotran, R.S., Reiss, C.S., Burakoff, S.J., Fiers, W., and Ault, K.A. (1983). Ia expression by vascular endothelium is inducible by activated T cells and by human gamma interferon. J Exp Med 157, 1339–1353.

Pober, J.S., Merola, J., Liu, R., and Manes, T.D. (2017). Antigen Presentation by Vascular Cells. Front Immunol 8, 1907.

Potente, M., and Makinen, T. (2017). Vascular heterogeneity and specialization in development and disease. Nat Rev Mol Cell Biol 18, 477–494.

Rothermel, A.L., Wang, Y., Schechner, J., Mook-Kanamori, B., Aird, W.C., Pober, J.S., Tellides, G., and Johnson, D.R. (2004). Endothelial cells present antigens in vivo. BMC Immunol 5, 5.

Rouhani, S.J., Eccles, J.D., Riccardi, P., Peske, J.D., Tewalt, E.F., Cohen, J.N., Liblau, R., Makinen, T., and Engelhard, V.H. (2015). Roles of lymphatic endothelial cells expressing peripheral tissue antigens in CD4 T-cell tolerance induction. Nat Commun 6, 6771.

Schaeuble, K., Cannelle, H., Favre, S., Huang, H.Y., Oberle, S.G., Speiser, D.E., Zehn, D., and Luther, S.A. (2019). Attenuation of chronic antiviral T-cell responses through constitutive COX2-dependent prostanoid synthesis by lymph node fibroblasts. PLoS Biol 17, e3000072.

Shaikh, H., Pezoldt, J., Mokhtari, Z., Gamboa Vargas, J., Le, D.D., Pena Mosca, J., Arellano Viera, E., Kern, M.A., Graf, C., Beyersdorf, N., et al. (2022). Fibroblastic reticular cells mitigate acute GvHD via MHCII-dependent maintenance of regulatory T cells. JCI Insight 7.

Shlomchik, W.D., Couzens, M.S., Tang, C.B., McNiff, J., Robert, M.E., Liu, J., Shlomchik, M.J., and Emerson, S.G. (1999). Prevention of graft versus host disease by inactivation of host antigen-presenting cells. Science 285, 412–415.

Sixt, M., Kanazawa, N., Selg, M., Samson, T., Roos, G., Reinhardt, D.P., Pabst, R., Lutz, M.B., and Sorokin, L. (2005). The conduit system transports soluble antigens from the afferent lymph to resident dendritic cells in the T cell area of the lymph node. Immunity 22, 19–29.

Tewalt, E.F., Cohen, J.N., Rouhani, S.J., Guidi, C.J., Qiao, H., Fahl, S.P., Conaway, M.R., Bender, T.P., Tung, K.S., Vella, A.T., et al. (2012). Lymphatic endothelial cells induce tolerance via PD-L1 and lack of costimulation leading to high-level PD-1 expression on CD8 T cells. Blood 120, 4772–4782.

Toubai, T., Tawara, I., Sun, Y., Liu, C., Nieves, E., Evers, R., Friedman, T., Korngold, R., and Reddy, P. (2012). Induction of acute GVHD by sex-mismatched H-Y antigens in the absence of functional radiosensitive host hematopoietic-derived antigen-presenting cells. Blood 119, 3844–3853.

Unanue, E.R. (2002). Perspective on antigen processing and presentation. Immunol Rev 185, 86–102.

Venneri, M.A., De Palma, M., Ponzoni, M., Pucci, F., Scielzo, C., Zonari, E., Mazzieri, R., Doglioni, C., and Naldini, L. (2007). Identification of proangiogenic TIE2-expressing monocytes (TEMs) in human peripheral blood and cancer. Blood 109, 5276–5285.

Warnock, R.A., Askari, S., Butcher, E.C., and von Andrian, U.H. (1998). Molecular mechanisms of lymphocyte homing to peripheral lymph nodes. J Exp Med 187, 205–216.

Welford, A.F., Biziato, D., Coffelt, S.B., Nucera, S., Fisher, M., Pucci, F., Di Serio, C., Naldini, L., De Palma, M., Tozer, G.M., and Lewis, C.E. (2011). TIE2-expressing macrophages limit the therapeutic efficacy of the vascular-disrupting agent combretastatin A4 phosphate in mice. J Clin Invest 121, 1969–1973.

Zeiser, R., and Blazar, B.R. (2017). Pathophysiology of Chronic Graft-versus-Host Disease and Therapeutic Targets. N Engl J Med 377, 2565–2579.

