## Supplementary material for "Secondary lymphoid organ endothelial cells prime alloreactive CD4^+^ T cells to trigger acute graft-versus-host disease": Main figures

### Slide 1
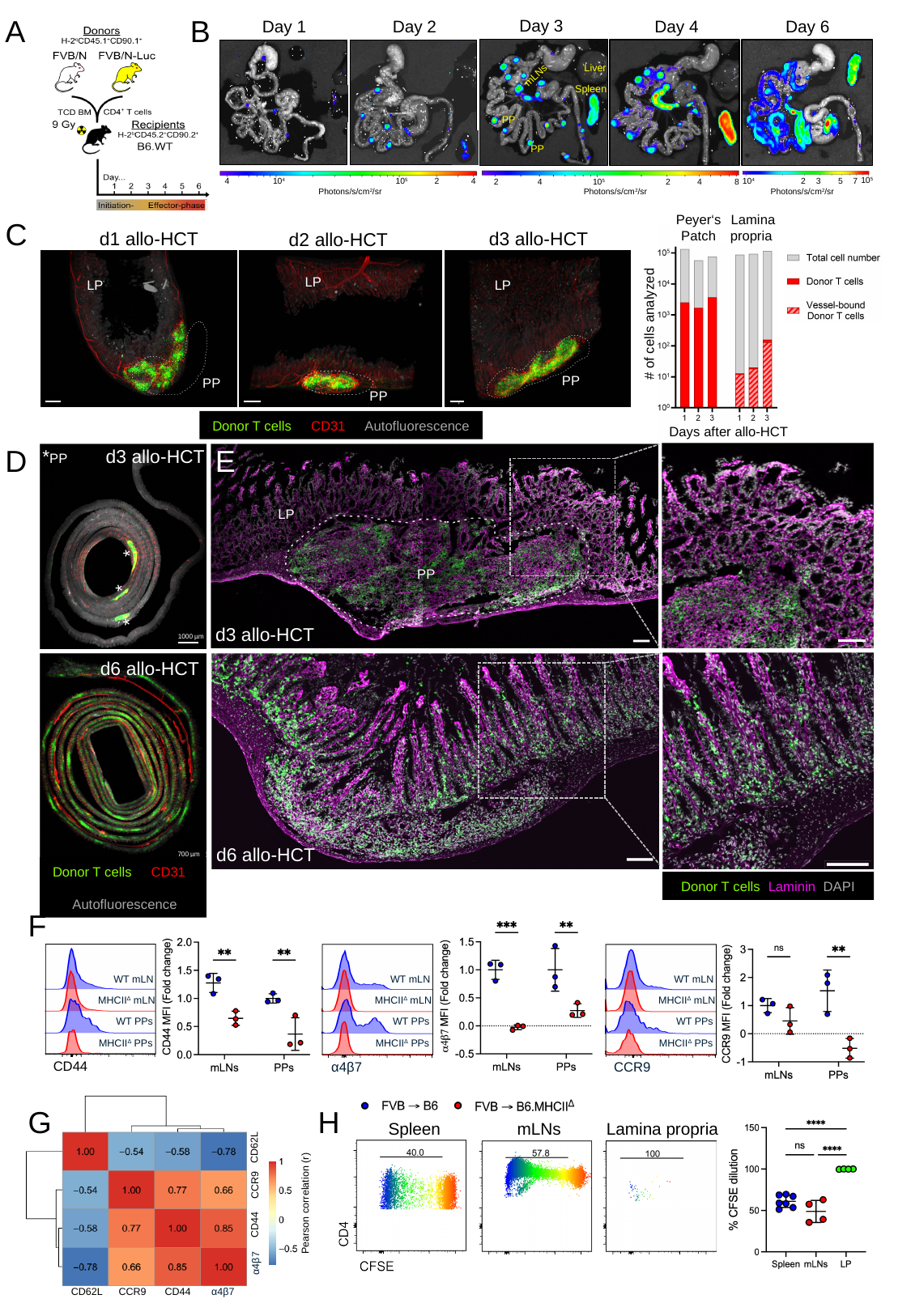

B
A
Day 3
Liver
mLNs
Spleen
PP
PP
Day 1
Day 2
Day 4
Day 6
4
10^4^
10^5^
2
4
Photons/s/cm^2^/sr
2
4
10^5^
2
4
8
Photons/s/cm^2^/sr
10^5^
5
10^4^
2
3
7
Photons/s/cm^2^/sr
Peyer‘s Patch
Lamina propria
C
d1 allo-HCT
d3 allo-HCT
d2 allo-HCT
LP
PP
LP
PP
LP
PP
#### of cells analyzed
50 µm
Donor T cells CD31 Autofluorescence
Days after allo-HCT
D
E
d3 allo-HCT
d6 allo-HCT
LP
PP
*PP
*
*
*
d3 allo-HCT
1000 µm
Donor T cells Laminin DAPI
d6 allo-HCT
700 µm
Donor T cells CD31
 Autofluorescence
Donor T cells Laminin DAPI
F
WT mLN
WT mLN
WT mLN
MHCII^Δ^ mLN
MHCII^Δ^ mLN
MHCII^Δ^ mLN
WT PPs
WT PPs
WT PPs
MHCII^Δ^ PPs
MHCII^Δ^ PPs
MHCII^Δ^ PPs
CD44
α4β7
CCR9
G
H
Lamina propria
mLNs
Spleen
CD62L
CCR9
Pearson correlation (r)
CD44
α4β7
CD62L
CCR9
CD44
α4β7

### Slide 2
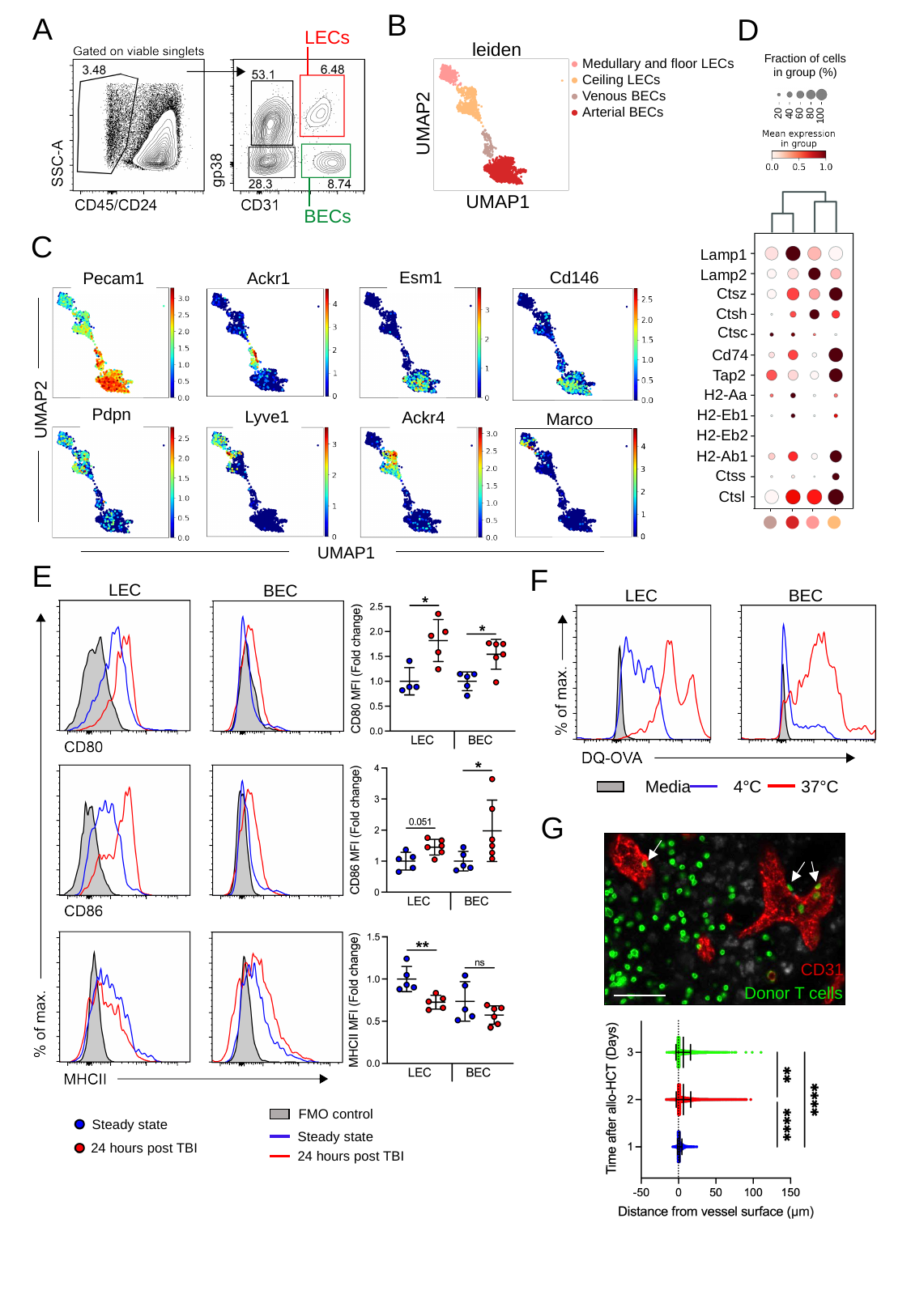

B
A
D
LECs
leiden
Fraction of cells
in group (%)
20
40
60
80
100
● Medullary and floor LECs
● Ceiling LECs
● Venous BECs
● Arterial BECs
UMAP2
Lamp1
Lamp2
Ctsz
Ctsh
Ctsc
Cd74
Tap2
H2-Aa
H2-Eb1
H2-Eb2
H2-Ab1
Ctss
Ctsl
●
●
●
●
UMAP1
BECs
C
Esm1
Cd146
Ackr1
Pecam1
UMAP2
Pdpn
Lyve1
Ackr4
UMAP1
Marco
E
F
LEC
BEC
LEC
BEC
37°C
Media
4°C
G
CD31
Donor T cells
FMO control
Steady state
24 hours post TBI
Steady state
24 hours post TBI

### Slide 3
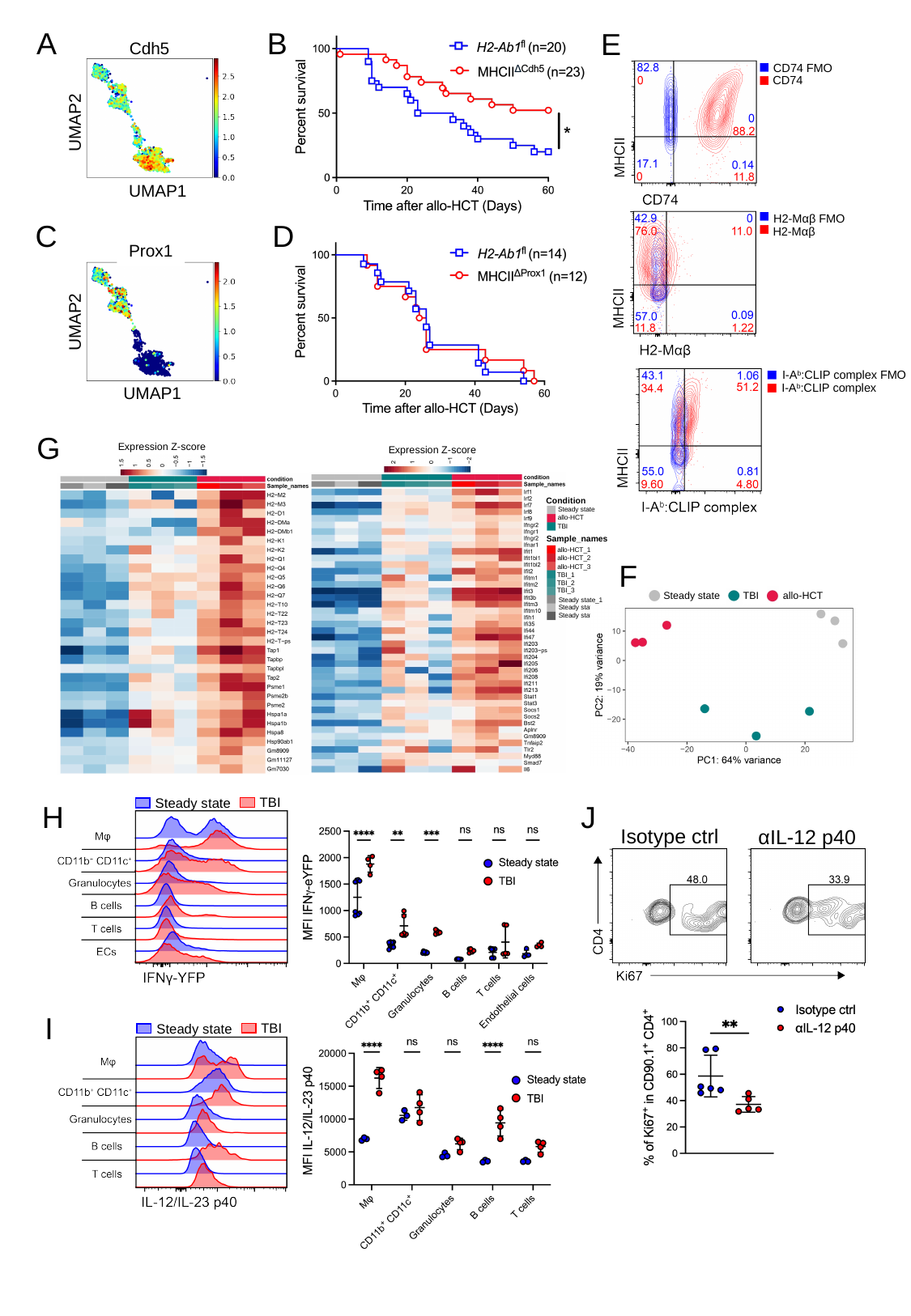

A
B
E
Cdh5
CD74 FMO
CD74
UMAP2
UMAP1
H2-Mαβ FMO
D
C
H2-Mαβ
Prox1
UMAP2
UMAP1
I-A^b^:CLIP complex FMO
I-A^b^:CLIP complex
G
Expression Z-score
Expression Z-score
Condition
Steady state
allo-HCT
TBI
Sample_names
allo-HCT_1
allo-HCT_2
allo-HCT_3
TBI_1
TBI_2
TBI_3
Steady state_1
Steady state_2
Steady state_3
F
Steady state
allo-HCT
TBI
TBI
Steady state
H
J
Isotype ctrl
αIL-12 p40
I
TBI
Steady state

### Slide 4
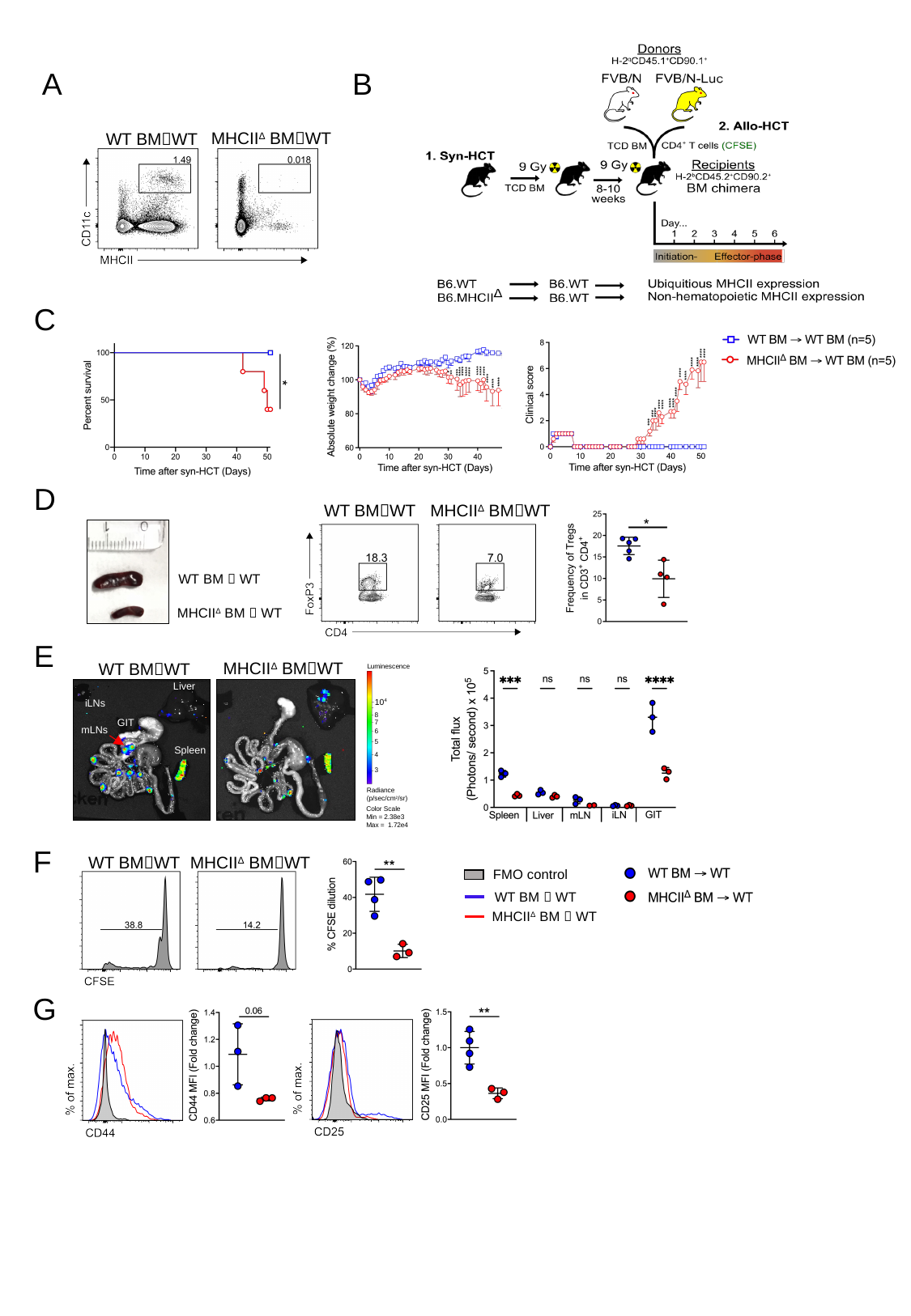

A
B
MHCII^Δ^ BM🡪WT
WT BM🡪WT
C
D
WT BM🡪WT
MHCII^Δ^ BM🡪WT
WT BM 🡪 WT
MHCII^Δ^ BM 🡪 WT
E
MHCII^Δ^ BM🡪WT
WT BM🡪WT
Luminescence
Liver
10^4^
iLNs
8
GIT
7
6
5
Spleen
4
3
Radiance
(p/sec/cm^2^/sr)
Color Scale
Min = 2.38e3
Max = 1.72e4
mLNs
F
WT BM🡪WT
MHCII^Δ^ BM🡪WT
FMO control
WT BM 🡪 WT
MHCII^Δ^ BM 🡪 WT
G

### Slide 5
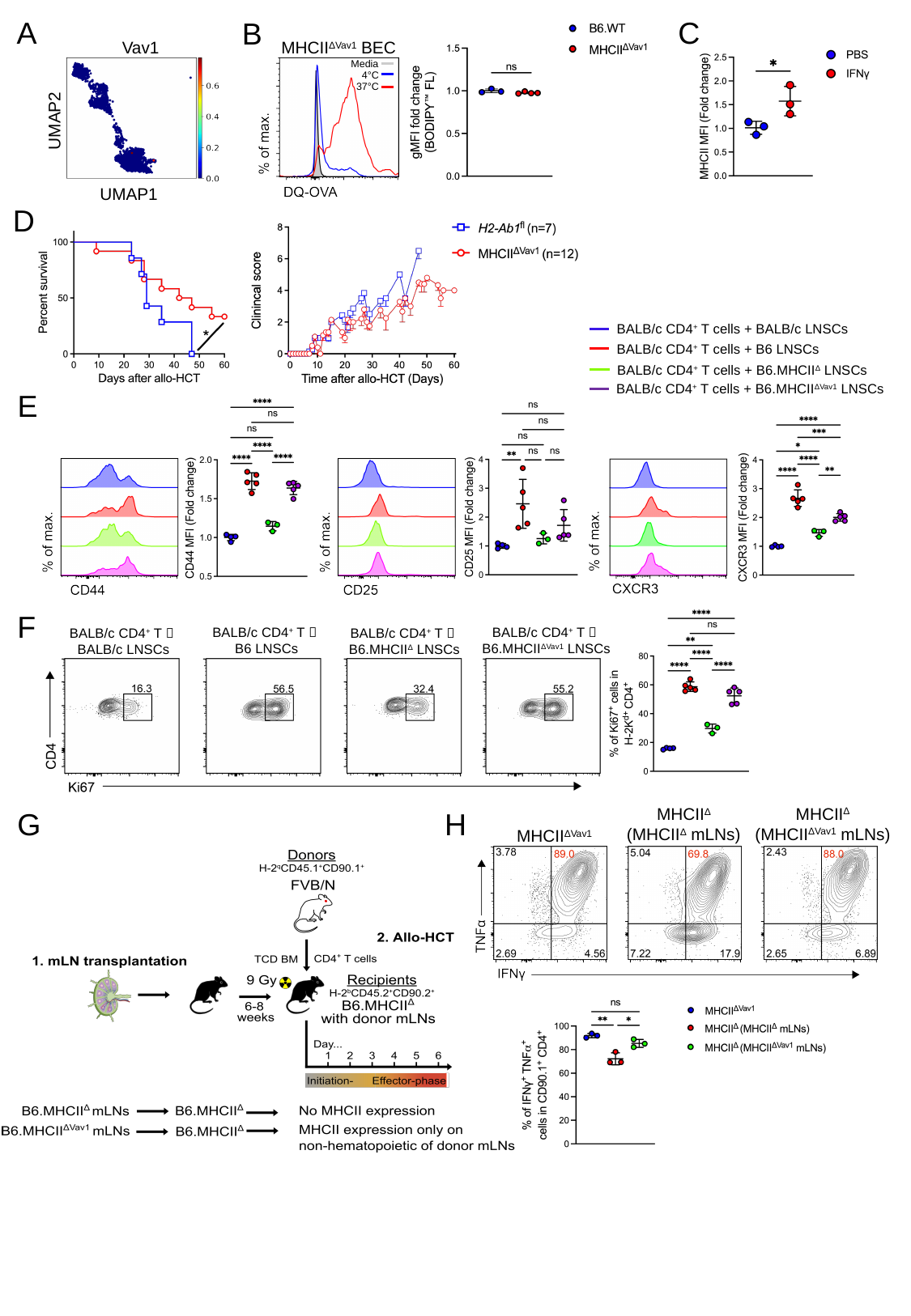

C
A
B
MHCII^ΔVav1^ BEC
Vav1
Media
4°C
37°C
UMAP2
UMAP1
D
BALB/c CD4^+^ T cells + BALB/c LNSCs
BALB/c CD4^+^ T cells + B6 LNSCs
BALB/c CD4^+^ T cells + B6.MHCII^ΔVav1^ LNSCs
BALB/c CD4^+^ T cells + B6.MHCII^Δ^ LNSCs
E
F
BALB/c CD4^+^ T 🡪
B6 LNSCs
BALB/c CD4^+^ T 🡪
B6.MHCII^ΔVav1^ LNSCs
BALB/c CD4^+^ T 🡪
BALB/c LNSCs
BALB/c CD4^+^ T 🡪
B6.MHCII^Δ^ LNSCs
MHCII^Δ^
(MHCII^ΔVav1^ mLNs)
MHCII^Δ^
(MHCII^Δ^ mLNs)
H
G
MHCII^ΔVav1^

### Slide 6
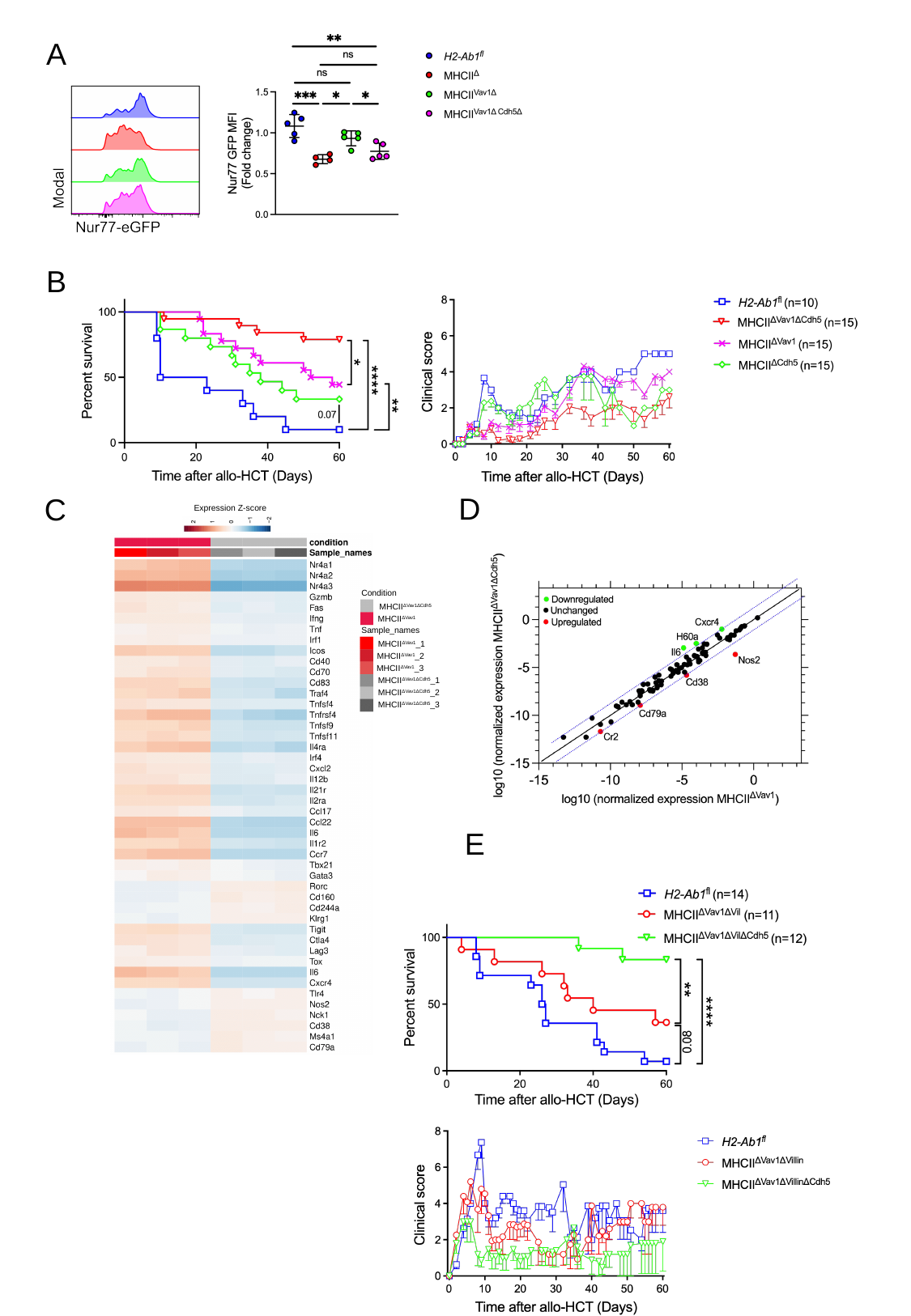

A
B
D
C
Expression Z-score
Condition
MHCII^ΔVav1ΔCdh5^
MHCII^ΔVav1^
Sample_names
MHCII^ΔVav1^_1
MHCII^ΔVav1^_2
MHCII^ΔVav1^_3
MHCII^ΔVav1ΔCdh5^_1
MHCII^ΔVav1ΔCdh5^_2
MHCII^ΔVav1ΔCdh5^_3
E
