## Supplementary figures for "Secondary lymphoid organ endothelial cells prime alloreactive CD4^+^ T cells to trigger acute graft-versus-host disease"

### Slide 1
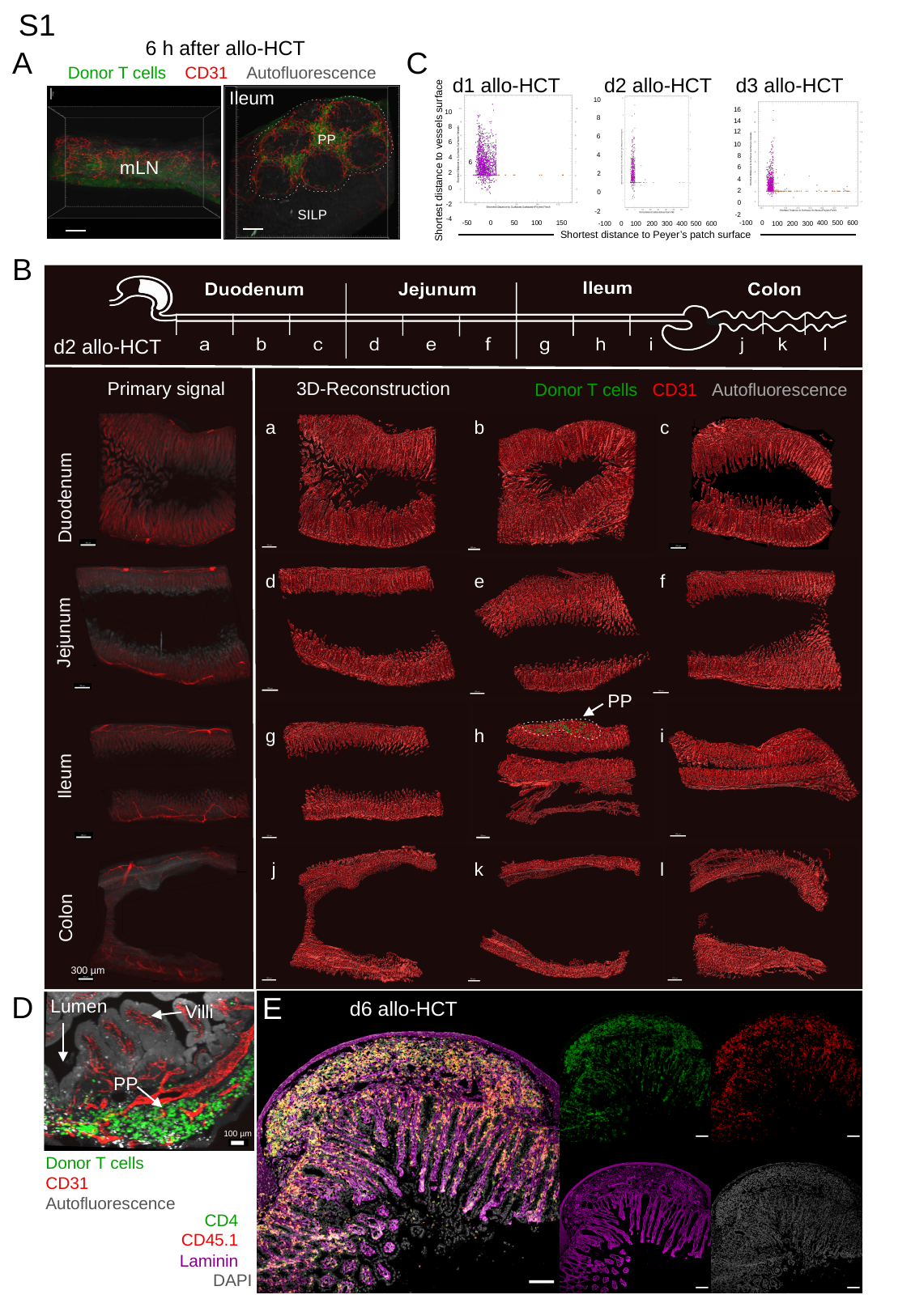

S1
6 h after allo-HCT
A
C
Donor T cells CD31 Autofluorescence
d1 allo-HCT
d2 allo-HCT
d3 allo-HCT
Ileum
PP
SILP
mLN
10
8
6
4
2
0
-2
16
14
12
10
8
6
4
2
0
-2
10
8
6
4
2
0
-2
-4
Shortest distance to vessels surface
6
-100
400
500
600
-50
0
50
100
150
0
200
300
100
500
-100
0
100
200
300
400
600
Shortest distance to Peyer’s patch surface
B
Duodenum
Jejunum
Ileum
Colon
300 µm
a
b
c
d
e
f
PP
g
h
i
j
k
l
Primary signal
3D-Reconstruction
Donor T cells CD31 Autofluorescence
d2 allo-HCT
D
E
d6 allo-HCT
Lumen
PP
100 µm
Villi
Donor T cells
CD31
Autofluorescence
CD4
CD45.1
Laminin
DAPI
Villus

### Slide 2
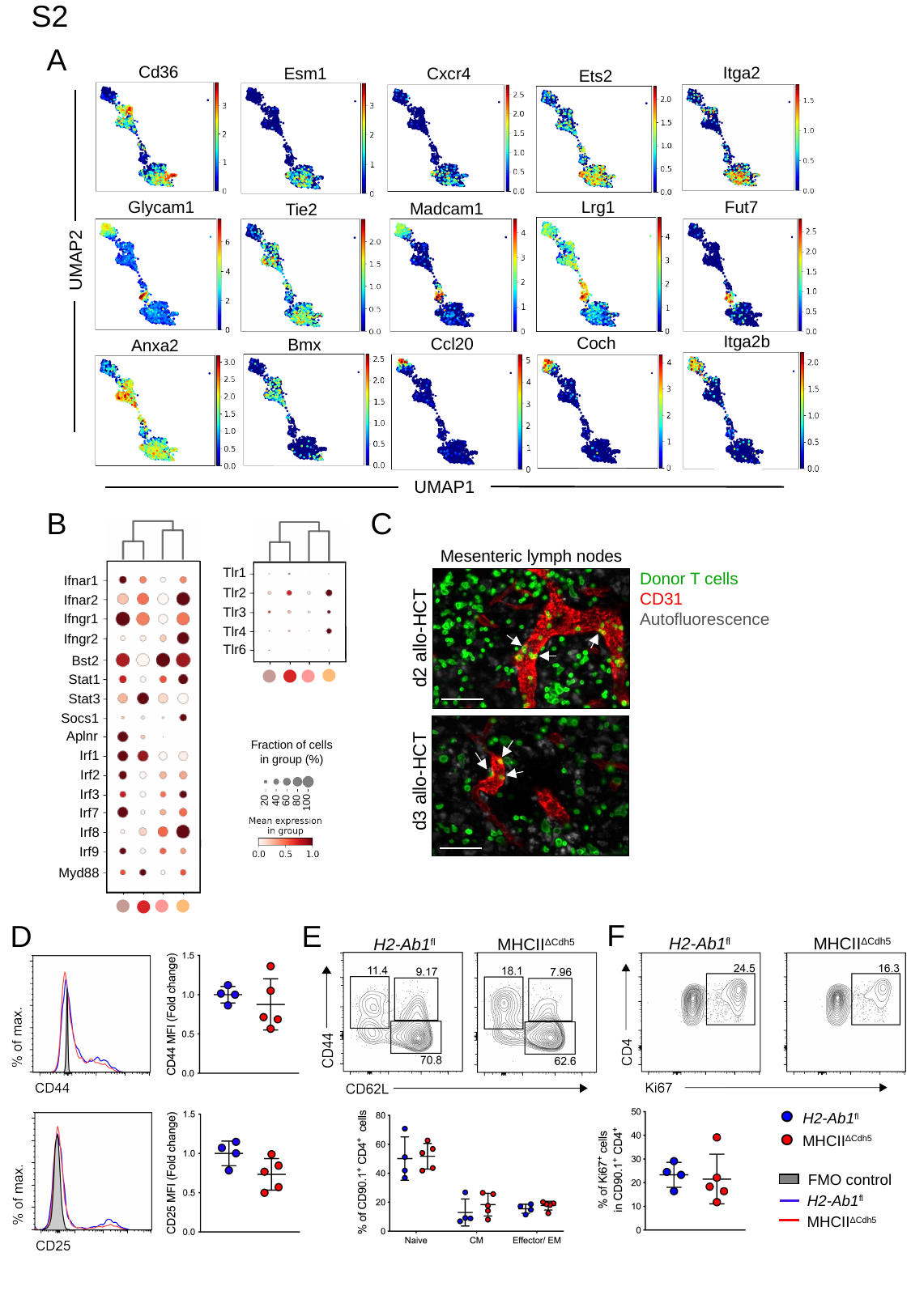

S2
A
Cd36
Itga2
Esm1
Cxcr4
Ets2
Lrg1
Glycam1
Fut7
Madcam1
Tie2
UMAP2
Itga2b
Coch
Ccl20
Bmx
Anxa2
UMAP1
B
C
Tlr1
Ifnar1
Tlr2
Ifnar2
Tlr3
Ifngr1
Tlr4
Ifngr2
Tlr6
Bst2
●
●
●
●
Stat1
Stat3
Socs1
Aplnr
Fraction of cells
in group (%)
20
40
60
80
100
Irf1
Irf2
Irf3
Irf7
Irf8
Irf9
Myd88
●
●
●
●
Mesenteric lymph nodes
Donor T cells
CD31
Autofluorescence
d2 allo-HCT
d3 allo-HCT
F
E
D
H2-Ab1^fl^
MHCII^ΔCdh5^
H2-Ab1^fl^
MHCII^ΔCdh5^
H2-Ab1^fl^
MHCII^ΔCdh5^
FMO control
H2-Ab1^fl^
MHCII^ΔCdh5^

### Slide 3
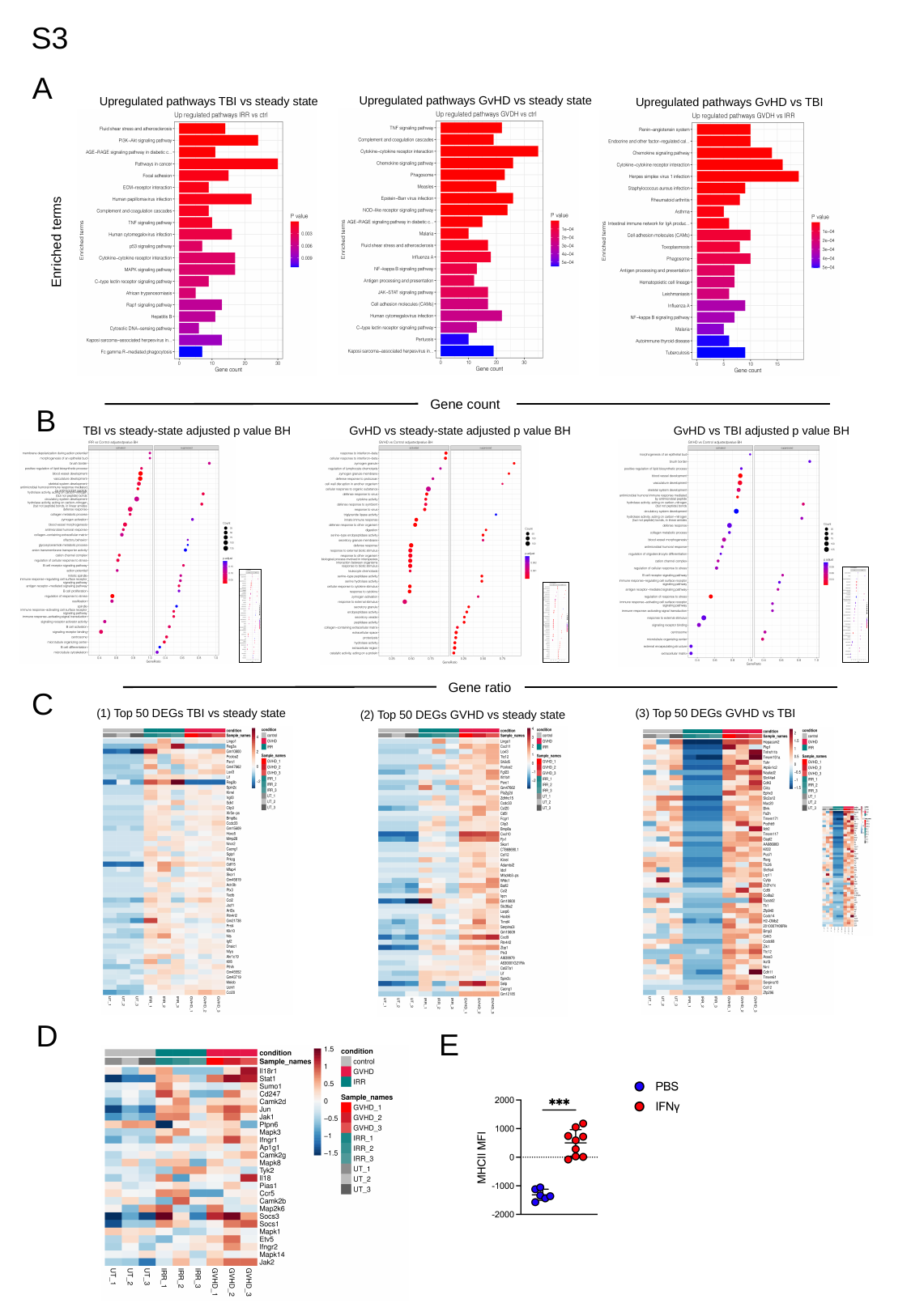

S3
A
Upregulated pathways GvHD vs steady state
Upregulated pathways TBI vs steady state
Upregulated pathways GvHD vs TBI
Enriched terms
Gene count
B
GvHD vs steady-state adjusted p value BH
TBI vs steady-state adjusted p value BH
GvHD vs TBI adjusted p value BH
Gene ratio
C
(1) Top 50 DEGs TBI vs steady state
(3) Top 50 DEGs GVHD vs TBI
(2) Top 50 DEGs GVHD vs steady state
D
E

### Slide 4
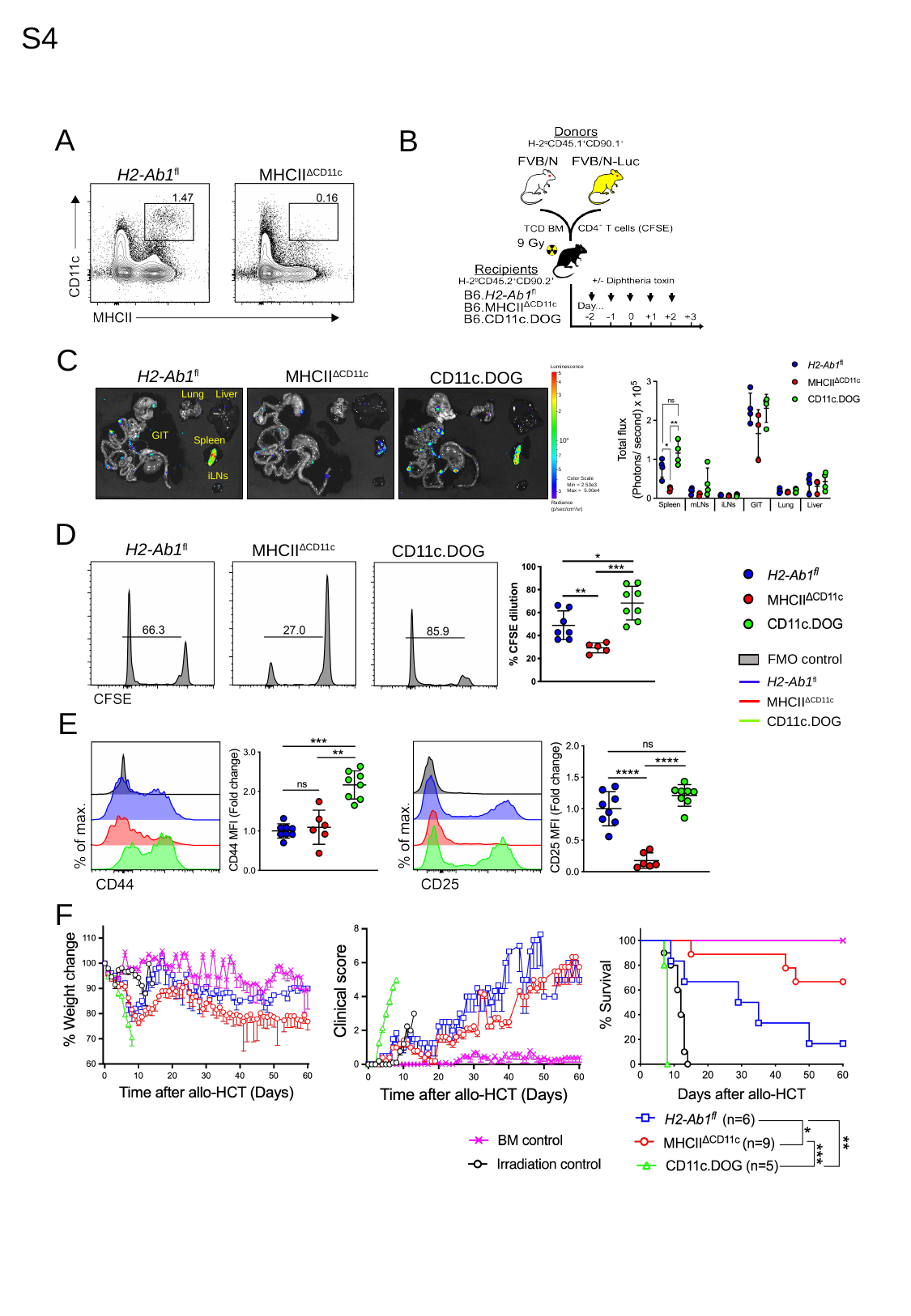

S4
A
B
H2-Ab1^fl^
MHCII^ΔCD11c^
C
Luminescence
5
4
3
2
10^4^
7
5
Color Scale
Min = 2.53e3
Max = 5.00e4
3
Radiance
(p/sec/cm^2^/sr)
H2-Ab1^fl^
MHCII^ΔCD11c^
CD11c.DOG
Lung
Liver
GIT
Spleen
iLNs
D
H2-Ab1^fl^
MHCII^ΔCD11c^
CD11c.DOG
FMO control
H2-Ab1^fl^
MHCII^ΔCD11c^
CD11c.DOG
E
F

### Slide 5
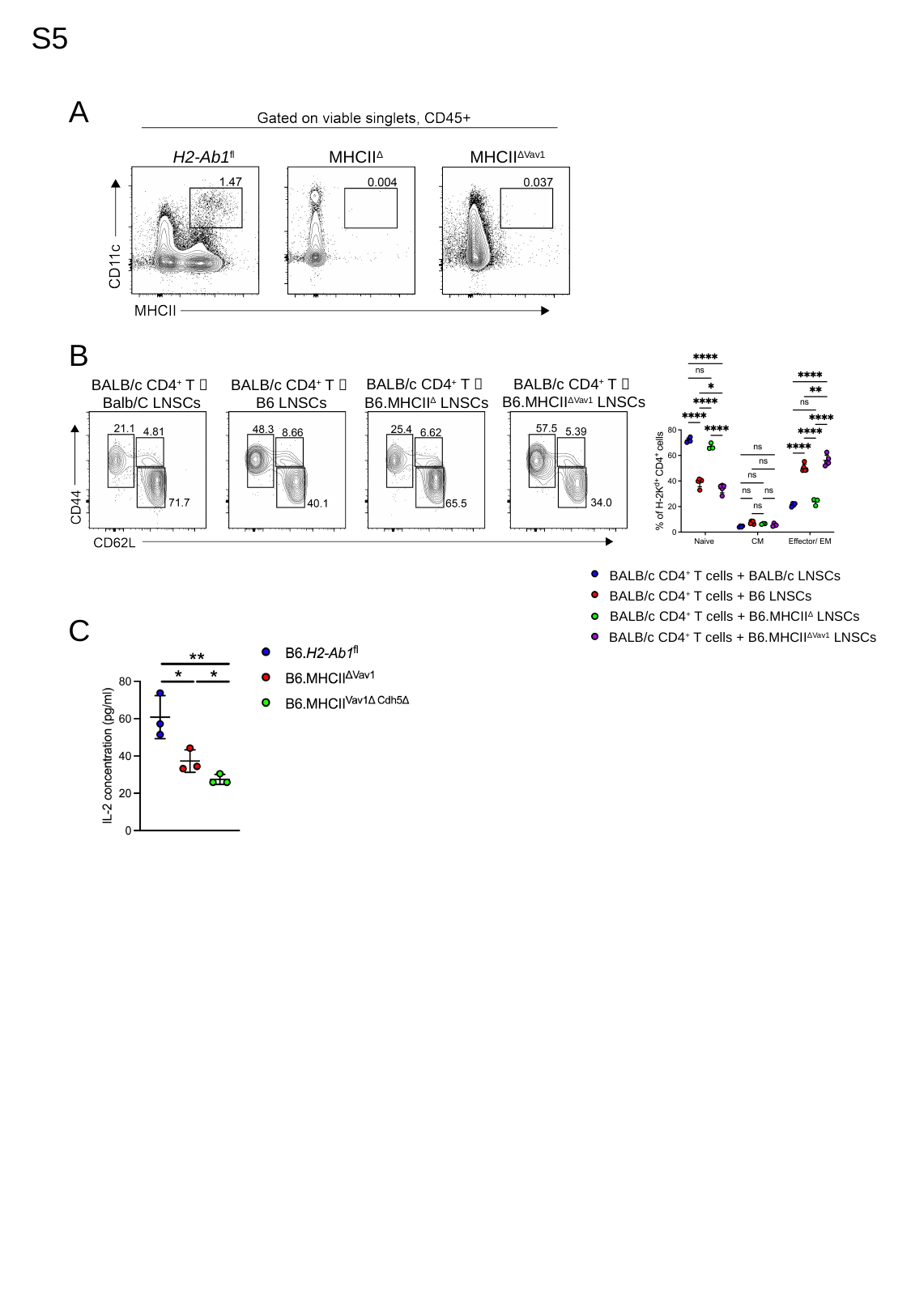

S5
A
MHCII^Δ^
MHCII^ΔVav1^
H2-Ab1^fl^
B
BALB/c CD4^+^ T 🡪
B6.MHCII^ΔVav1^ LNSCs
BALB/c CD4^+^ T 🡪
B6.MHCII^Δ^ LNSCs
BALB/c CD4^+^ T 🡪
Balb/C LNSCs
BALB/c CD4^+^ T 🡪
B6 LNSCs
BALB/c CD4^+^ T cells + BALB/c LNSCs
BALB/c CD4^+^ T cells + B6 LNSCs
BALB/c CD4^+^ T cells + B6.MHCII^Δ^ LNSCs
BALB/c CD4^+^ T cells + B6.MHCII^ΔVav1^ LNSCs
C

### Slide 6
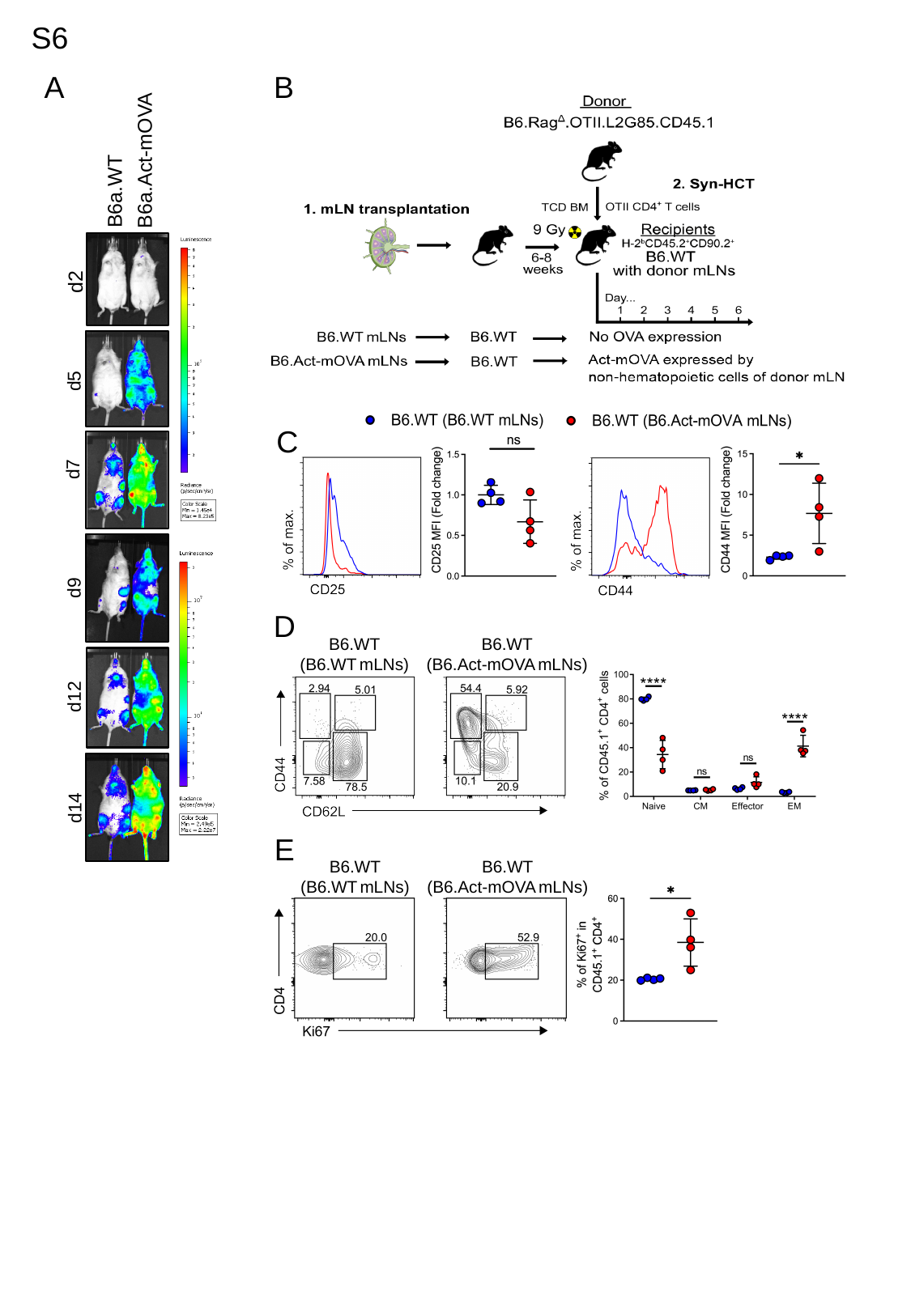

S6
B
A
B6a.Act-mOVA
B6a.WT
d2
d5
d7
d9
d12
d14
C
D
B6.WT
(B6.WT mLNs)
B6.WT
(B6.Act-mOVA mLNs)
E
B6.WT
(B6.WT mLNs)
B6.WT
(B6.Act-mOVA mLNs)

### Slide 7
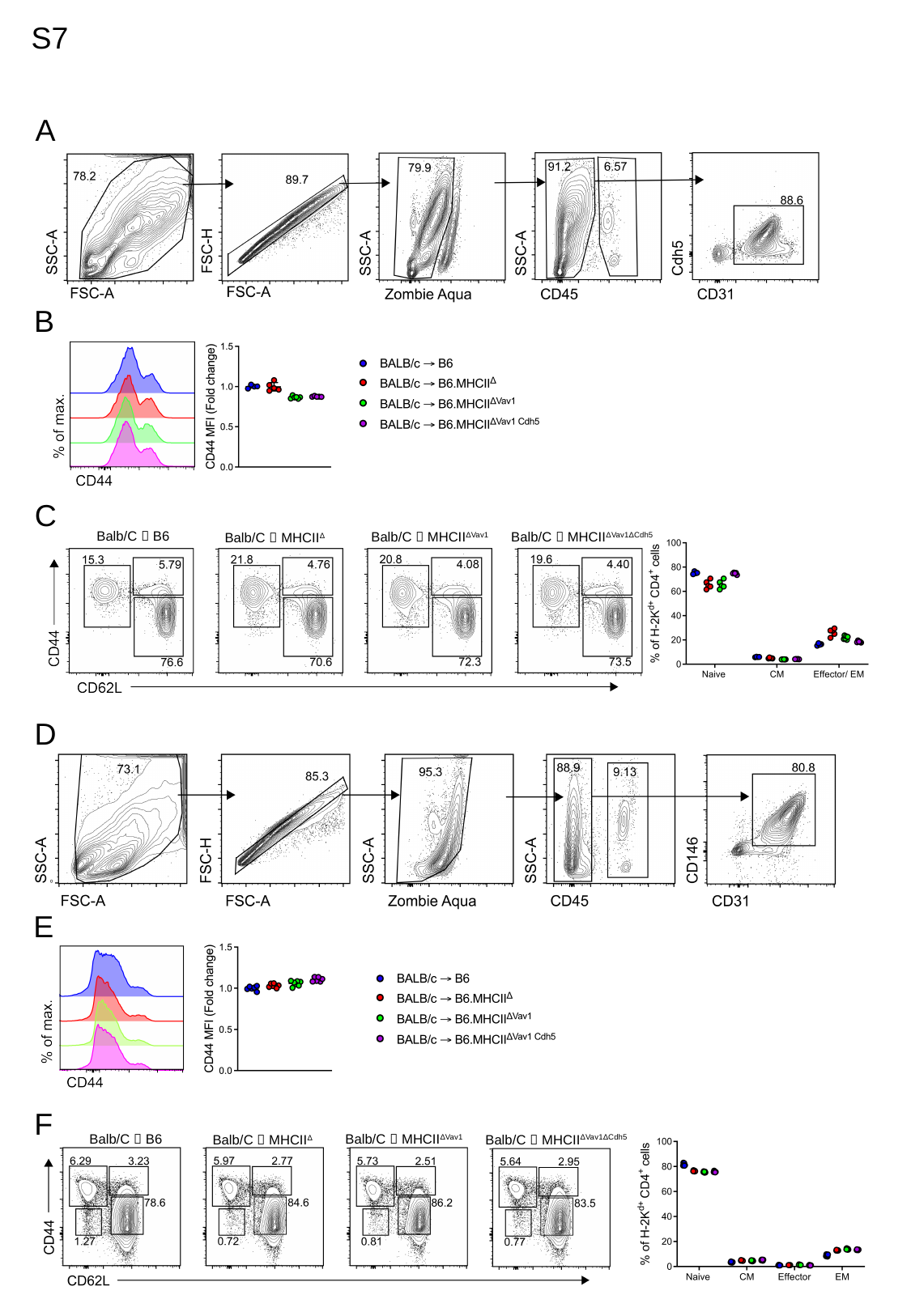

S7
A
B
C
Balb/C 🡪 B6
Balb/C 🡪 MHCII^ΔVav1^
Balb/C 🡪 MHCII^ΔVav1ΔCdh5^
Balb/C 🡪 MHCII^Δ^
D
E
F
Balb/C 🡪 B6
Balb/C 🡪 MHCII^ΔVav1^
Balb/C 🡪 MHCII^ΔVav1ΔCdh5^
Balb/C 🡪 MHCII^Δ^
