## Supplementary material for "Secondary lymphoid organ endothelial cells prime alloreactive CD4^+^ T cells to trigger acute graft-versus-host disease": Material and methods

**Mice.** C57BL/6J (B6.WT, H-2^b^, CD45.2), FVB (FVB, H2-2^q^, CD45.1), and BALB/c (C, H-2^d^ CD45.2) mice were obtained from Charles River Laboratories (Sulzfeld, Germany) and Janvier Laboratories (Le Genest-Saint-Isle, France). C57BL/6-background CD11c-Cre, Vav1-iCre, Cdh5-Cre, Villin-Cre-ER^T2^, *H2-Ab1*^fl^, GREAT (IFNg-eYFP), Act-mOVA and MHCII^Δ^ (null) mice were obtained from The Jackson Laboratories (Bar Harbor, Maine, USA). B6.CD11c.DOG mice (Hochweller et al.) were provided by Günter J. Hämmerling (German Cancer Research Center, Heidelberg, Germany). *H2-Ab1*^fl^ mice were crossed with CD11c-Cre, Vav1-iCre, Cdh5-Cre, Prox1-Cre and Villin-Cre mice to generate MHCII^ΔCD11c^, MHCII^ΔVav1^, MHCII^ΔCdh5^, MHCII^ΔProx1^ and MHCII^ΔVil^ mice, respectively. Subsequently, MHCII^ΔVav1^ and MHCII^ΔCdh5^ were crossed to generate animals MHCII-deficient in hematopoietic and endothelial l cells. MHCII^ΔVav1^ and MHCII^ΔVil^ were crossed to generate MHCII-deficient mice in hematopoietic and intestinal epithelial cells. MHCII^ΔVav1^, MHCII^ΔVil^ and MHCII^ΔCdh5^ were crossed to generate MHCII-deficient mice in hematopoietic, endothelial and intestinal epithelial cells. BALB/c.Nur77-eGFP (C, H-2^d^, CD45.2) (Moran et al.) were kindly provided by Kristin A. Hogquist (University of Minnesota, United States). FVB/L2G85 (H-2^q^, CD90.1, CD45.1) (Beilhack et al.; Cao et al.) were bred inhouse. B6.Rag^Δ^.OTII.L2G85.CD45.1 were generated by crossing Rag^Δ^ mice (Mombaerts et al.) with mice expressing OVA-specific TCR on CD4^+^ T cells (Barnden et al.) crossed to B6 mice harbouring the CAG-luc-eGFP L2G85 transgene carrying *Ptprc*^a^ (Bauerlein et al.; Cao et al.). All mice were maintained under specific pathogen-free (SPF) conditions in individually ventilated cages (IVCs) at the Center for Experimental Molecular Medicine (ZEMM), University of Würzburg, Germany.

**METHOD DETAILS**

**Mixed lymphocyte reaction (MLR).** MLRs were performed using responder CD4^+^ T cells enriched from the spleens of FVB/N or C.Nur77-eGFP mice using Dynabeads Untouched Mouse CD4^+^ Cells Kit (Invitrogen), following manufacturer’s instructions. Stimulator cells were obtained from LNs following enzymatic digestion (Fletcher et al.) and subsequent depletion of hematopoietic cells using MojoSort Mouse CD45 Nanobeads (Biolegend). Co-cultures were maintained in complete RPMI (cRPMI) supplemented with 1% penicillin-streptomycin, 1% insulin-transferrin-selenium (ITS) solution (Gibco), 10% fetal calf serum (FCS) (Gibco), IL-2 (100 U/ml) (Biolegend), IL-7 (10 ng/ml) (Biolegend), and incubated for 5 days at 37°C in a humidified CO₂ incubator.

For the isolation of endothelial cells, mice were intracardially perfused with DPBS and an enzymatic mix containing collagenase D and DNase I (Sigma Aldrich) for liver processing (Sokol et al.). Lung tissue was processed using the Mouse Lung Dissociation Kit (Miltenyi Biotec) following perfusion with DPBS. Endothelial cells were enriched via positive selection using CD146 (LSEC) and CD31 MicroBeads (Miltenyi Biotec). Isolated endothelial cells were then co-cultured with allogeneic CD4^+^ T cells in cRPMI supplemented with IL-2 (100 U/ml), IL-7 (10 ng/ml), endothelial cell growth supplement/heparin (1.2 μg/ml) and Normocin™ (100 μg/ml), and incubated for 5 days at 37°C in a humidified CO₂ incubator.

**Bone marrow transplantation.** Sex-matched (B6, H-2^b^) recipient mice aged 8- to 12-weeks were subjected to myeloablative total body irradiation (TBI) with a single dose of 9 Gy (Faxitron CP-160 x ray irradiation system, Faxitron X-Ray). Within 4 hours after TBI, mice were i.v. injected with 5x10^6^ T cell–depleted (TCD) BM cells from either FVB or BALB/c donors for hematopoietic reconstitution. T cells were depleted from BM using CD90.1 MicroBeads (Miltenyi Biotec) for FVB and Dynabead^TM^ Mouse Pan T (Thy1.2) kit (Invitrogen) for BALB/c, following manufacturers’ instructions. To induce aGvHD, allogeneic CD4^+^ T cells enriched from FVB, FVB.L2G85 or C.Nur77-GFP mice were co-injected intravenously. CD4^+^ T cells were isolated using Dynabeads Untouched Mouse CD4^+^ Cells Kit (Invitrogen) according to manufacturer’s instructions. Cell purity was confirmed by flow cytometry and consistently exceeded 95% purity. For IL-12 blockade, mice received intraperitoneal injections of 500 µg of anti-IL-12p40 (C17.8) or isotype control rat IgG2a (2A3) antibody from day -3 to +2 relative to allo-HCT.

**Bone marrow chimeras.** B6.WT recipient mice were pre-treated for one week with acidified drinking water (pH 2.5-3) supplemented with 1 mg/ml gentamycin sulphate solution (Sigma Aldrich) prior to irradiation (9 Gy) and i.v. injection of 1x10^7^ TCD BM in 200 µl DPBS from either B6.WT (B6.WT 🡪 B6.WT) or MHCII^Δ^ (B6.MHCII^Δ^ 🡪 B6.WT) for hematopoietic reconstitution. T cells were depleted using the Dynabead^TM^ Mouse Pan T (Thy1.2) kit (Invitrogen).

Eight to twelve weeks post-reconstitution, chimerism was assessed by flow cytometry, and only mice with confirmed engraftment were used for subsequent allo-HCT experiments.

**Mesenteric lymph node transplantation.** Lymph node transplantation surgery was performed as previously described (Shaikh et al.). Briefly, mice were anesthetized, shaved, and administered an intradermal subcutaneous (s.c.) analgesic injection (Carprofen, Midas Pharma GmbH). A 1-1.5 cm midline incision was made through the skin and peritoneum along the linea alba. Sterile swabs were used to gently open the peritoneum cavity and exteriorize the caecum and small intestine with minimal handling to avoid post-operative ileus. Using fine surgical scissors, the recipient’s mLNs mice were excised, ensuring minimal removal of surrounding adipose tissue. Donor mLNs were then transplanted into the mesenteric tissue using fibrin glue (TISSEEL, Baxter). 20 µl fibronectin was applied to the excised recipient mLN area, and donor mLNs–pre-dipped in thrombin solution–were placed in the correct anatomical orientation (from distal to the proximal). The glue was allowed to set for 3-5 min, after which the small and large intestine, along with the caecum, were gently repositioned into the peritoneal cavity. The peritoneum was closed with three non-continuous sutures using coated VICRYL® suture (polyglactin 910, Ethicon), and the skin was closed with four individual stiches. Bepanthen® Wund- und Heilsalbe (Bayer) was applied to the external wound. Mice were placed in a clean recovery cage and kept warm under an infrared lamp until they had fully recovered from anesthesia.

Transplanted mice were allowed to rest 5-10 weeks to ensure revascularization and integration of the donor LN, including angiogenesis and lymphatic reconnection.

**Flow cytometry.** *in* *vitro*–cultured cells and primary mouse single-cell suspensions were incubated with normal rat serum (NRS) (NRS, 1:20 DPBS) for 5 minutes at 4°C to block non-specific Fc receptor binding. Cells were stained with fluorochrome-conjugated antibodies for 30 minutes at 4°C. Dead cells were excluded using LIVE/DEAD Fixable Violet Dead Cell Stain Kit (Invitrogen) or Zombie Aqua™ Fixable Viability Kit (Biolegend). Data was acquired on a BD FACS Canto II flow cytometer (BD Biosciences) or Attune NxT flow cytometer (Thermo Fisher Scientific), and analyzed using FlowJo version 10 software (BD Biosciences). UltraComp eBeads Compensation Beads (Invitrogen) were stained with individual fluorochrome–labelled antibodies for compensation, allowing correction of spectral overlap between fluorophores. Fluorescence minus one (FMO) controls (Herzenberg et al.) were used to define gates in multicolor panels. For fluorescence activated cell sorting (FACS), cells were antibody-stained and sorted using a FACS Aria III (BD Biosciences) with a 100 μm nozzle into ice-cold cell culture medium or lysis buffer depending on downstream application. CD4^+^ T cells, LECs and BECs were sorted to >95% purity.

**Antigen processing assay.** FACS-sorted cells (>95% purity) were incubated with 10 μg/ml BODIPY-conjugated DQ-OVA (Invitrogen) in RPMI-1640 supplemented with 1% penicillin-streptomycin and 10% FCS at 4°C and 37°C for 3 hours. DQ-OVA processing was measured by flow cytometry on an Attune NxT flow cytometer under the blue laser (488 nm, BL-1-channel).

**RT^2^ Profiler PCR Array.**

CD4^+^ T cells were enriched from spleen and LNs of B6.MHCII^ΔVav1^ and B6.MHCII^ΔVav1ΔCdh5^ mice at day 22 of allo-HCT. RNA was extracted using the RNeasy Mini Kit (Qiagen) and converted to cDNA with the RT^2^ First Strand Kit (Qiagen). Real time PCR was performed using the RT^2^ SYBR Green qPCR mastermix (Qiagen) on Mouse T-cell & B-cell activation array (Bio-Rad CFX96). C_T_ values were normalized with reference genes (*Actb*, *B2m*, *Gapdh*, *Hprt1*, *Rplp0*) using the 2-^ΔΔCt^ method, and data were visualized as a scatter plot.

**Single-cell RNA-Seq data analysis.** Publicly available single-cell RNA-Seq data (GEO: GSE116633; (Pezoldt et al.) were analyzed using SCANPY (Wolf et al.) in Python. Cells with <500 or >5,000 >7.5% mitochondrial reads were filtered out (Ilicic et al.). Genes detected in fewer than 3 cells were excluded. Counts were log-normalized, and 3,000 variable genes were selected. Total counts (<https://www.cureffi.org/2013/09/12/counts-vs-fpkms-in-rnaseq/>) and mitochondrial gene percentage were regressed out, and data were scaled to unit variance.

Dimensional reduction was performed using principal component analysis (PCA, 40 components) and embedded in 2D using UMAP (20 neighbours). Clustering was done via Leiden algorithm (resolution = 1). Endothelial cells were identified by *Pecam* (CD31) expression > 1.5, and 2,013 cells from 2 mLNs and 2 peripheral LNs were re-clustered. Clusters were annotated using dot plots of marker genes and differentially expressed genes (DEGs).

**RNA-Seq and analysis.** Myeloablative conditioned B6.WT mice received 5x10^6^ TCD BM and 6x10^5^ CD4^+^ T cells from FVB donors. On day 3.5 after allo-HCT, LNs were digested and CD45^+^ cells were depleted CD45 MicroBeads (Miltenyi Biotec) according to manufacturer’s instructions. The enriched CD45^-^ cell fraction was stained with LIVE/DEAD Fixable Violet Dead Cell Stain Kit (Invitrogen) to exclude non-viable cells, and surface markers CD45, CD24, CD31 and gp38 to identify ECs. Viable CD45^-^CD24^-^gp38^-^CD31^+^ BECs were sorted using a 100 μm nozzle in 150 μl of lysis buffer and stored on dry ice for downstream processing.

For CD4^+^ T cell RNA seq, spleens and LNs were collected 22 days after allo-HCT, processed into single-cell suspension, and stained with LIVE/DEAD Fixable Violet Dead Cell Stain Kit (Invitrogen). Donor-derived CD4^+^ T cells were identified by staining for CD90.1 and CD4, and sorted using a 70 μm nozzle in 150 μl of lysis buffer, then stored on dry ice.

Total RNA was extracted using the PicoPure RNA isolation kit (Thermo Fisher Scientific) following manufacturer’s protocol. RNA integrity and concentration was assessed using a 2100 Bioanalyzer instrument (Aligent). cDNA synthesis and library preparation were performed using 0.5 ng of RNA with the NEBNext Single Cell/Low Input RNA Library Prep kit for Illumina, according to the manufacturer’s instructions. Libraries were sequenced on an Illumina NextSeq 500 system with 75 bp single-end reads. Read quality was assessed using FastQC.

Fastq files were processed using the R package “Rsubread” (v2.14.0) for alignment and gene-level read counting (Liao et al.), using the Genome Reference Consortium Mouse Build 38 (GRCm38) reference genome. Raw count matrices were imported into “DESeq2” (v 1.40.0) for downstream analysis (Love et al.). Genes with fewer than 10 reads and detected in fewer than 3 samples were excluded as part of quality control. PCA was performed on vst-transformed data, using the top 500 most variable genes. Differentially expressed genes (DEGs) between conditions were computed using “DESeq” function. Volcano plots were generated using shrunken log_2_ fold change (LFC) and standard errors estimated with “apeglm” (Zhu et al.). Visualization was performed using the "pheatmap", “EnhancedVolcano”, “RColorBrewer” and “ggvenn” R packages. For gene set enrichment analysis (GSEA), DEGs with log_2_ fold change between -2 and +2 were ranked and submitted to gseGO or gseKEGG functions from the clusterProfiler R package, using a p-value cutoff of 0.1 and Benjamini-Hochberg (BH) adjustment for multiple testing (Wu et al.). RNA-Seq data have been deposited in the Gene Expression Omnibus (GEO) under accession numbers GSE267745 (for BECs) and GSE267746 (for CD4^+^ T cells).

**IL-2 assay.** IL-2 secretion by BO-97.10 hybridomas (Hugo et al.) was measured after 5 days of co-culture with the CD45^-^ fraction isolated from LNs of *H2-Ab1*^fl^, MHCII^Δ^, MHCII^ΔVav1^, and MHCII^ΔCdh5ΔVav1^ mice. Following co-culture, IL-2 concentrations in the culture supernatant were quantified using a Mouse IL-2 ELISA Kit (Thermo Fisher Scientific) according to the manufacturer’s instructions.

**BLI.** In *vivo* BLI was performed using the IVIS Spectrum CCD-imaging system (PerkinElmer) as previously described (Chopra et al.). Mice were anesthetized via i.p.-injected of a ketamine (50 μg/g body weight) and xylazine (5 μg/g body weight) mixture in DPBS, with a total injection volume of 10 μl/g body weight. Mice received an i.p. injection of D-Luciferin (300 mg/kg body weight), and images were acquired 10 minutes after injection to assess T cell proliferation and migration. Alternatively, mice were anesthetized using 2% isoflurane in O_2_ and injected i.p. with 300 mg/kg of D-Luciferin. After 10 minutes, bioluminescence signals were acquired using the IVIS Spectrum (PerkinElmer).

For *ex vivo* imaging, mice were injected with the same anesthetic and D-Luciferin regimen. After 10 minutes, mice were euthanized, and organs were harvested within 4 minutes post-euthanasia for imaging. *Ex vivo* BLI enabled enhanced resolution of bioluminescent signal distribution across specific organs. Imaging analysis was conducted using Living image 4.5.5 software (PerkinElmer).

**Immunofluorescence staining.** Mice were perfused intravascularly with DPBS for 2 minutes, followed by 4% PFA for 8 minutes to ensure tissue fixation. Dissected tissues were further post-fixed in 4% PFA for 3 h at room temperature. Following fixation, tissues were cryoprotected by sequential equilibration in 10% sucrose overnight at 4°C, 20% sucrose for 4 h, and finally 30% sucrose for 2 h. Cryoprotected tissues were embedded in OCT compound and sectioned at 7 μm thickness using a cryostat (CM1900; Leica Biosystems). Tissue sections were mounted on frosted glass slides and blocked for 30 minutes in 2% fetal calf serum (FCS) in DPBS, using an avidin/biotin blocking kit (Vector Laboratories) to minimize background staining. Slides were incubated with primary antibodies (see Key Resources Table) for 1 h at room temperature, followed by incubation with the appropriate fluorophore-conjugated secondary antibodies for 30 minutes. Nuclei were counterstained with DAPI, and slides were mounted using mounting medium (Vector Laboratories). Images were acquired at room temperature using a confocal laser-scanning microscope (LSM780; ZEISS) and analyzed with IMARIS software version 8.1.1 (Bitplane AG).

**3D light sheet fluorescence microscopy (LSFM) staining and analysis.** Three-dimensional LSFM staining was performed as previously described (Brede et al.; Shaikh et al.). Briefly, primary antibodies were administered intravenously 2 h prior to perfusion, followed by secondary antibody injection 30 minutes before intracardiac perfusion. Mice were first perfused with ice-cold DPBS for 2 minutes, followed by 4% PFA in DPBS for 8 minutes. Harvested organs were post-fixed in 4% PFA for 3 hours at 4°C, then washed three times in cold DPBS (15 minutes each). Samples were permeabilized by overnight incubation in 0.1% Triton X-100 in DPBS at 4°C. Dehydration was carried out at room temperature through a graded ethanol series (30%, 50%, 70%, 80%, 90%, and 100%) for 90 minutes per step, with samples left in 100% ethanol at 4°C overnight. For optical clearing, ethanol was replaced with n-hexane for 2 hours at room temperature. Subsequently, n-hexane was gradually replaced with BABB (1 part benzyl alcohol, 2 parts benzyl benzoate) in three steps, carefully removing solution from the top while simultaneously adding BABB to ensure specimens remained submerged. Finally, the clearing solution was replaced with fresh BABB and samples were incubated overnight at 4°C.

To generate 3D Swiss Rolls of the ileum (Mueller et al.) the tissue was opened longitudinally along the mesenteric border and transferred into a flat dish containing PBS. Using fine forceps, the specimen was carefully rolled—starting from the proximal end—with the luminal side facing outward around a wooden swab (LP Italiana, 112298; cotton tips removed). The swab was then placed into a large processing cassette (Macrosette M512, Simport), trimmed to fit, and immediately fixed in 4% PFA. Following fixation, the rolled tissues were gently unravelled in ice-cold PBS to remove residual chyme and faeces, and re-rolled in the same orientation onto plastic stirring rods (Brand, VWR 441-0217). Rolls were transferred back into cassettes for dehydration and clearing, as described above. After the final dehydration step, plastic rods were removed prior to immersion in BABB.

3D-LSFM imaging was performed using a custom-built LSFM system optimized for whole-organ imaging, equipped with four excitation wavelengths (491, 532, 642, and 730 nm), as previously described (Brede et al.). Multicolor image stacks were acquired in 2 µm or 5 µm steps, with sequential capture across all color channels. The system was controlled by IQ 2.9 software (Andor, Belfast, UK), and image data were saved in TIFF format. Image processing and quantitative analysis were conducted using IMARIS software (v10.2.0; Bitplane AG, Switzerland). When necessary, background subtraction was applied based on the estimated diameter of the targeted cell population to reduce non-specific signals. Cellular and vascular structures were reconstructed using the “Surface” tool, applying a smoothing factor of 0.5 µm for cells and 1 µm for vessels, with manually defined intensity thresholds. To quantify the spatial relationship between donor T cells and PPs in the small intestine, an iso-surface was manually created by outlining the boundaries of PPs in each Z-stack slice. The “Distance Transformation” and “Find Nearest Neighbors” functions were used to calculate the distance between T cells and either PPs surfaces or mLNs vasculature.

**Statistics.** Data are shown as mean ± SD. Comparisons between two groups were performed using two-tailed unpaired Student’s t test or the non-parametric Mann–Whitney U test, as appropriate. For comparisons involving more than two groups, two-way ANOVA was used, followed by Tukey’s multiple comparisons test for post hoc analysis. Survival data were analyzed using Kaplan–Meier survival curves, and statistical differences were assessed with the log-rank (Mantel–Cox) test. A *p*-value of less than 0.05 was considered statistically significant. All statistical analyses were conducted using GraphPad Prism 10 (GraphPad Software, San Diego, CA).

**Materials availability.** This study did not generate new unique reagents.

**Data and code availability.** scRNA-seq data has been deposited at GEO and is publicly available as of the date of publication. Accession numbers are listed in the key resource table. This paper does not report original code. Any additional information required to re-analyze the data reported in this paper is available from the lead contact upon request.

**Materials**

| **Antibodies** | | |
| --- | --- | --- |
| Alexa Fluor 647 anti-mouse CD11c (clone N418) | BioLegend | Cat. # 117312; RRID: AB_389328 |
| APC/Cyanine7 anti-mouse/human CD11b (clone M1/70) | BioLegend | Cat. # 101226; RRID: AB_830642 |
| PerCP/Cyanine5.5 anti-mouse CD24 (clone M1/69) | BioLegend | Cat. # 101824; RRID: AB_1595491 |
| PerCP/Cyanine5.5 anti-mouse TER-119 (clone TER-119) | BioLegend | Cat. # 116228; RRID: AB_893636 |
| APC anti-mouse CD25 (clone PC61) | BioLegend | Cat. # 102012; RRID: AB_312861 |
| Alexa Fluor 488 anti-mouse CD31 (clone MEC13.3) | BioLegend | Cat. # 102514; RRID: AB_2161031 |
| Biotin anti-mouse CD31 (clone 390) | BioLegend | Cat. # 102404; RRID: AB_312899 |
| PE-Cyanine7 anti mouse CD3ε (clone 145-2C11) | Invitrogen | Cat. # 25-0031-82; RRID: AB_469572 |
| PerCP/Cyanine5.5 anti-mouse CD4 (clone RM4-5) | BioLegend | Cat. # 100540; RRID: AB_893326 |
| PE anti-mouse/human CD44 (clone IM7) | BioLegend | Cat. # 103008; RRID: AB_312959 |
| Pacific Blue anti-mouse CD45 (clone s30-F11) | Invitrogen | Cat. # MCD4528; RRID: AB_10373710 |
| APC/Cyanine7 anti-mouse CD45.1(clone A20) | BioLegend | Cat. # 110716; RRID: AB_313505 |
| PE/Cyanine7 anti-mouse CD62L (clone MEL-14) | BioLegend | Cat. # 104418; RRID: AB_313505 |
| PE/Cyanine7 anti-mouse CD80 (clone 16-10A1) | BioLegend | Cat. # 104734; RRID: AB_2563113 |
| APC/Cyanine7 anti-mouse CD86 (clone GL1) | BioLegend | Cat. # 105030; RRID: AB_2244452 |
| APC-eFluor 780 anti-mouse CD90.1 (clone HIS51) | eBioscience | Cat. # 47-0900-82; RRID: AB_1272252 |
| Biotin anti-mouse CD90.1 (clone HIS51) | eBioscience | Cat. # 13-0900-81; RRID: AB_466530 |
| PerCP-Cyanine5.5 anti mouse CXCR3 (clone CXCR3-173) | Invitrogen | Cat. # 45-1831-82; RRID: AB_1210699 |
| APC anti-mouse FoxP3 (clone FJK-16s) | eBioscience | Cat. # 17-5773-82; RRID: AB_469457 |
| PE anti-mouse I-A^b^ (clone  AF6-120.1) | BioLegend | Cat. # 116408; RRID: AB_313727 |
| Alexa Fluor 647 anti-mouse Ki-67 (clone 16A8) | BioLegend | Cat. # 652408; RRID: AB_2562139 |
| APC anti-mouse Podoplanin (clone 8.1.1) | BioLegend | Cat. # 127410; RRID: AB_10613649 |
| APC-eFluor™ 780 anti-mouse H-2K^d^ (clone SF1-1.1.1) | eBioscience | Cat. # 47-5957-82; RRID: AB_2762719 |
| Alexa Fluor 647 Donkey anti-rabbit IgG | BioLegend | Cat. # 406414; RRID: AB_2563202 |
| PE anti-mouse TNFα (clone MP6-XT22) | BD Pharmingen | Cat. # 554419; RRID: AB_395380 |
| Alexa Fluor 488 anti-mouse IFNγ (clone XMG1.2) | eBioscience | Cat. # 53-7311-82; RRID: AB_469932 |
| Alexa Fluor® 647 anti-mouse Cdh5 (clone BV13) | BioLegend | Cat. # 138006; RRID: AB_10569114 |
| PerCP/Cyanine5.5 anti-mouse CD146 (clone ME-9F1) | BioLegend | Cat. # 134710; RRID: AB_11203708 |
| PE anti-mouse/human B220 (clone  RA3-6B2) | BioLegend | Cat. # 103208; RRID: AB_312993 |
| APC anti-mouse α4β7 (clone LPAM-1) | Invitrogen | Cat. # 17-5887-82; RRID: AB_1210577 |
| PE-Cy7 anti-mouse CCR9 (clone CW-1.2) | Invitrogen | Cat. # 25-1991-82; RRID: AB_10854423 |
| PE anti-mouse IL-12/IL-23 p40 (clone C17.8) | eBioscience | Cat. # 12-7123-82; RRID: AB_466185 |
| Anti-mouse IL-12 p40 (clone C17.8) | BioXCell | Cat. # BE0051; RRID: AB_1107698 |
| Rat IgG2a isotype control, anti-trinitrophenol (clone 2A3) | BioXCell | Cat. # BE0089; RRID: AB_1107769 |
| Alexa Fluor 647 anti-mouse CLIP (clone In1) | BioLegend | Cat. # BE0051; RRID: AB_2632608 |
| PE anti mouse MHC class II (clone 15G4) | Santa Cruz Biotechnology | Cat. # sc-53946 PE; RRID: N/A |
| Purified rat anti-mouse H2-M (clone 2E5A) | BD Pharmingen | Cat. # 552405; RRID: AB_394380 |
| Purified rabbit anti-mouse Laminin (polyclonal) | Invitrogen | Cat. # PA1-16730; RRID: AB_2133633 |
| Biotin anti-mouse CD45.1 (clone A20) | eBioscience | Cat. # 13-0453-82; RRID: AB_466453 |
| Purified Rat anti-mouse CD4 (clone H129.19) | BD Biosciences | Cat. # 550278; RRID: AB_394967 |
| Rat anti-Mouse IgG1 Secondary Antibody, APC | eBioscience | Cat. # 17-4015-82; RRID: AB_2573205 |
| Cy™3 AffiniPure® Fab Fragment Donkey Anti-Rabbit IgG (H+L) | Jackson ImmunoResearch | Cat. # 711-167-003; RRID: AB_2340606 |
| Donkey anti-Rat IgG (H+L) Antibody, Alexa Fluor™ Plus 488 | Invitrogen | Cat. # A48269; RRID: AB_2893137 |
| **Chemical, peptides, and recombinant proteins** | | |
| Benzyl benzoate | Sigma Aldrich | Cat. # B6630-1L |
| Benzyl alcohol | Sigma Aldrich | Cat. # 305197-2L |
| Collagenase D | Roche | Cat. # 11088858001 |
| DPBS | PAN Biotec | Cat. # P04-36500 |
| DNase I | Sigma Aldrich | Cat. # DN25 |
| Dispase II | Roche | Cat. # 4942078001 |
| Corn oil | Sigma Aldrich | Cat. # C8267 |
| RPMI 1640 Medium | Gibco | Cat. # 11875093 |
| Fetal Bovine Serum | Sigma Aldrich | Cat. # F7524 |
| Endothelial Cell Growth Supplement / Heparin | PromoCell | Cat. # C-30120 |
| PMA/Ionomycin (500X) | Biolegend | Cat. # 423302 |
| Insulin-Transferrin-Selenium (ITS -G) (100X) | Thermo Fischer Scientific | Cat. # 41400045 |
| RT2 First Strand Kit | Qiagen | Cat. # 330401 |
| RT² SYBR Green qPCR Mastermix | Qiagen | Cat. # 330500 |
| Gentamycin Sulphate | Sigma Aldrich | Cat. # BP174 |
| Brefeldin A Solution (1000X) | BioLegend | Cat. # 420601 |
| D-Luciferin | Biosynth | Cat. # L-8220 |
| Recombinant mouse IL-2 | BioLegend | Cat. # 75406 |
| Recombinant mouse IL-7 | BioLegend | Cat. # 577804 |
| RNeasy Mini Kit | Qiagen | Cat. # 74104 |
| PicoPure RNA Isolation Kit | Thermo Fischer Scientific | Cat. # KIT0204 |
| IL-2 mouse ELISA kit | Thermo Fischer Scientific | Cat. # BMS601 |
| TISSEEL | Baxter | Cat. # 021899-41891 |
| Hanks’ Balanced Salt solution (HBSS) | Gibco | Cat. # 14175-129 |
| Ketamine | Pfizer | Cat. # N/A |
| LIVE/DEAD Fixable Violet Dead Cell Stain | Thermo Fischer Scientific | Cat. # L34964 |
| UltraComp eBeads™ Compensation Beads | Thermo Fischer Scientific | Cat. # 01-2222-41 |
| Xylazine | CP Pharma | N/A |
| Zombie Aqua™ Fixable Viability Kit | BioLegend | Cat. # 423102 |
| Streptavidin, Alexa Fluor™ 647 Conjugate | Thermo Fischer Scientific | Cat. # S21374 |
| **Experimental Models: Organism/Strains** | | |
| Mouse: B6.WT: C57BL/6JRj | Jackson laboratories | RRID: IMSR_JAX:000664 |
| Mouse: B6a.WT: C57BL/6JRj-Tyr^c/c^ | Jackson laboratories | RRID: IMSR_JAX:000058 |
| Mouse: BALB/c: BALB/cAnNRj | Charles River | RRID: IMSR_CRL:028 |
| Mouse: FVB: FVB/NRj | Jackson laboratories | RRID: IMSR_JAX:001800 |
| Mouse: CD11c.DOG: *B6.Tg(Itgax-DTR/OVA/EGFP)1Garbi* | Dr. Günter J. Hammerling | Hochweller et al., 2008 |
| Mouse: C.Nur77-eGFP: C-*Tg(Nr4a1-EGFP/cre)820Khog* | Dr. Kristin A. Hogquist | Moran et al., 2011 |
| Mouse: FVB.L2G85: FVB-*Tg(CAG-luc,-GFP)L2G85Chco/*J | Dr. Robert Negrin | Cao et al., 2004 |
| Mouse: CD11c-Cre: B6.Cg-*Tg(Itgax-cre)1-1Reiz/*J | Jackson laboratories | RRID: IMSR_JAX:008068 |
| Mouse: Vav1-iCre: B6.Cg-*Tg(VAV1-cre)1Graf/MdfJ* | Jackson laboratories | RRID: IMSR_JAX:008610 |
| Mouse: Cdh5-iCre: B6.FVB-*Tg(Cdh5-cre)7Mlia*/J | Jackson laboratories | RRID: IMSR_JAX:006137 |
| Mouse: Villin-Cre: B6.SJL-*Tg(Vil-cre)997Gum*/J | Jackson laboratories | RRID: IMSR_JAX:004586 |
| Mouse: Prox1-Cre: Prox1tm3(cre/ERT2)Gco/J | Jackson laboratories | RRID: IMSR_JAX:022075 |
| Mouse: H2-Ab1^fl^: B6.129X1-*H2-Ab1tm1Koni*/J | Jackson laboratories | RRID: IMSR_JAX:013181 |
| Mouse: MHCII^Δ^: B6.129S2-*H2dlAb1-Ea*/J | Jackson laboratories | RRID: IMSR_JAX:003584 |
| Mouse: B6.129S4-*Ifng^tm3.1Lky^*/J | Jackson laboratories | RRID: IMSR_JAX:017581 |
| Mouse: Act-mOVA: B6-*Tg(CAG-OVAL)916Jen*/J | Jackson laboratories | RRID: IMSR_JAX:005145 |
| Mouse: B6.Rag^Δ^.OTII.L2G85.CD45.1 | ZEMM, University of Würzburg | Barnden et al., 1998; Mombaerts et al., 1992; Cao et al., 2004; Bäuerlein et al., 2013 |
| Each of the Cre strain and *H2-Ab1*^fl^ were inter-crossed to generate: |  |  |
| Mouse: MHCII^ΔCD11c^ | ZEMM, University of Würzburg | N/A |
| Mouse: MHCII^ΔCdh5^ | ZEMM, University of Würzburg | N/A |
| Mouse: MHCII^ΔVav1^ | ZEMM, University of Würzburg | N/A |
| Mouse: MHCII^ΔProx1^ | ZEMM, University of Würzburg | N/A |
| Mouse: MHCII^ΔVil^ | ZEMM, University of Würzburg | N/A |
| Mouse: MHCII^ΔVav1ΔCdh5^ | ZEMM, University of Würzburg | N/A |
| Mouse: MHCII^ΔVav1ΔVil^ | ZEMM, University of Würzburg | N/A |
| Mouse: MHCII ^ΔVav1ΔCdh5ΔVil^ | ZEMM, University of Würzburg | N/A |
| **Critical commercial Assays** | | |
| eBioscience Foxp3 / Transcription Factor Staining Buffer Set | eBioscience | Cat. # 00-5523-00 |
| Dynabeads™ Untouched™ Mouse CD4 Cells Kit | Thermo Fischer Scientific | Cat. # 11415D |
| Dynabeads™ Mouse Pan T (Thy1.2) | Thermo Fischer Scientific | Cat. # 11443D |
| CD90.1 MicroBeads | Miltenyi Biotec | Cat. # 130-121-273 |
| CD45 MicroBeads | Miltenyi Biotec | Cat. # 130-052-301 |
| CD146 (LSEC) MicroBeads | Miltenyi Biotec | Cat. # 130-092-007 |
| CD31 MicroBeads | Miltenyi Biotec | Cat. # 130-097-418 |
| MojoSort™ Mouse CD45 Nanobeads | BioLegend | Cat. # 480028 |
| RT^2^ PCR profiler | Qiagen | Cat. # 330231 |
| Lung Dissociation Kit, mouse | Miltenyi Biotec | Cat. # 130-095-927 |
| DQ^TM^ Ovalbumin | Thermo Fischer Scientific | Cat. # D-12053 |
| **Deposited Data** | | |
| Bulk RNA sequencing raw data | This paper | GEO: GSE267745, GEO: GSE267746 |
| Single cell RNA sequencing raw data | Pezoldt et al. 2018 | GEO: GSE116633 |
| **Software and Algorithms** | | |
| BD FACSDiva software version 8 | BD Biosciences | RRID: SCR_001456 |
| FlowJo v10 | BD Biosciences | RRID: SCR_008520 |
| GraphPad Prism (ver. 7.02) | GraphPad Software | RRID: SCR_002798 |
| IMARIS v8.3 | Bitplane AG | RRID: SCR_007370 |
| ImageJ | NIH | RRID: SCR_003070 |
| GSEA | Broad Institute | RRID: SCR_003199 |
